# A TERRA–NONO Axis Drives Fibroblast Reprogramming in Cancer

**DOI:** 10.64898/2026.06.08.730872

**Authors:** Emery Di Cicco, Jovan Isma, Stefano Sol, Gaetana Restivo, Atul Katarkar, Sandro Goruppi, Joni Renee White, Juerg Hafner, Victor A. Neel, Anna Mandinova, Mitchell P Levesque, Gian Paolo Dotto

## Abstract

The molecular mechanisms linking chromosome homeostasis to fibroblast reprogramming into cancer-associated fibroblasts (CAFs) remain poorly understood. Here, we identify a clinically relevant, telomere-associated nuclear pathway that drives CAF activation across multiple human cancers. The long non-coding RNA TERRA is consistently elevated in CAFs from skin, lung, and breast tumors. We show that the androgen receptor (AR) normally represses TERRA transcription by binding subtelomeric regions. Loss of AR releases this repression, allowing TERRA to interact with the RNA-binding protein NONO, forming a CAF-specific nuclear complex that reprograms transcription toward a tumor-promoting state. Disrupting the TERRA–NONO complex, through TERRA silencing or pharmacologic NONO inhibition, reverses CAF activation and suppresses tumor–stroma interactions both in vitro and in vivo. Importantly, NONO inhibition restores normal fibroblast features in patient-derived actinic keratoses, cutaneous squamous cell carcinomas, and melanomas, underscoring the translational potential of this pathway and positioning the TERRA–NONO complex as a promising therapeutic target.

## Introduction

The rising incidence of cancer with age is driven by converging biological processes, including telomere alterations at chromosome ends and epigenetic reprogramming within both cancer cells and their microenvironment (*1*). Whether these hallmarks are functionally interconnected remains an open question.

The long non-coding RNA TERRA (telomeric repeat-containing RNA) is a critical regulator of telomere function. It modulates telomeric chromatin architecture, influences telomerase activity, and participates in the alternative lengthening of telomeres (ALT) pathway (*2*). Beyond telomeres, TERRA associates with internal chromosomal loci and interacts with chromatin remodelers and transcriptional regulators, suggesting its role in broader epigenetic control (*3*) (*4*).

Among TERRA’s binding partners is NONO (also known as p54nrb), a multifunctional member of the DBHS (Drosophila behavior/human splicing) protein family. NONO functions as a molecular scaffold in processes ranging from transcription and RNA processing to genome maintenance (*5*). At telomeres, NONO binds TERRA and preserves their integrity by suppressing the formation of harmful RNA: DNA R-loops (*6*). However, the role of the TERRA–NONO complex beyond telomeres remains poorly defined. TERRA transcripts range from 100 base pairs to over 9 kilobases and originate from subtelomeric regions across multiple chromosomes (*2*). TERRA has been implicated in telomere length control in either a positive or negative fashion in connection with telomerase activity (*7*, *8*). Its expression varies significantly across various cancer cell types, being both upregulated and downregulated, suggesting context-dependent functions (*9*). Whether or not TERRA plays an intrinsic role in stromal fibroblasts and their activation into cancer-associated fibroblasts (CAFs), key stromal players that drive tumor progression (*10–12*), has not been explored.

Environmental damaging agents such as UVA radiation and cigarette smoke can penetrate the stromal compartment and induce long-lasting pro-tumorigenic changes. In human skin, downregulation of the Androgen Receptor (AR) has been shown to promote senescence and conversion of human dermal fibroblasts (HDFs) into CAFs (*13*) (*14*).

Here, we demonstrate that TERRA, a long non-coding RNA transcribed from multiple subtelomeric regions, is under direct negative regulation by AR in HDFs and is consistently upregulated in CAFs across major skin cancer types as well as in lung and breast cancer. We find that TERRA functions as both a necessary and sufficient driver of CAF transcriptional reprogramming. Quantitative proteomic analysis identifies the RNA-binding protein NONO as a key effector of TERRA function. Silencing NONO recapitulates the effects of TERRA depletion in reversing the CAF gene expression program, while pharmacological disruption of the TERRA–NONO complex abrogates CAF activation and suppresses tumorigenesis. Together, these findings uncover a previously unrecognized role for TERRA–NONO complex in governing stromal plasticity and reveal a therapeutically targetable mechanism of CAF reprogramming in the tumor microenvironment.

## RESULTS

### TERRA is upregulated in CAFs across multiple tumor types

The expression and role of the telomeric long non-coding RNA TERRA in the tumor stroma remain largely undefined. TERRA is transcribed from the subtelomeric regions of most human chromosomes (*2*). Using RT-qPCR with subtelomeric primers (1q, 2q, 7p, 9q, 13q, 17p, XqYq) we detected consistently elevated TERRA levels in six CAF strains derived from cutaneous squamous cell carcinomas (SCCs) compared to matched human dermal fibroblasts (HDFs) from adjacent skin, with the highest level of TERRA transcription from chromosome 7p (Fig.1A, fig. S1A, B). TERRA upregulation correlated with increased expression of canonical CAF markers, as previously reported (*13*, *14*). Increased TERRA levels were also seen in CAFs from basal cell carcinomas and melanomas when compared to reference HDFs from non-diseased areas (Fig.1B; Fig. S1C). These findings were validated by RNA-FISH using a probe targeting the repetitive sequence of TERRA, which revealed a marked increase in nuclear TERRA foci in CAFs compared to matched HDFs (Fig.1C, Fig.S1D). To validate these findings in patient specimens, immuno-RNA FISH performed on five human cSCC biopsies revealed significantly increased TERRA accumulation in vimentin-positive stromal fibroblasts relative to adjacent normal skin (Fig. 1D, Fig.S1E). Notably, this pattern was recapitulated in a tissue microarray of 37 primary lung SCC and adenocarcinoma samples, where vimentin-positive stromal cells displayed significantly elevated TERRA foci compared to matched normal lung tissue (Fig. 1E, Fig.S1F). A parallel analysis of breast cancer tissue microarray showed a striking upregulation of TERRA in stromal fibroblasts (Fig. 1F, Fig. S1G). Collectively, these findings establish TERRA accumulation in CAFs as a conserved hallmark of the tumor stroma in skin, lung, and breast cancers, underscoring its potential as a broadly applicable biomarker and mechanistic driver of stromal reprogramming.

**Figure 1.**
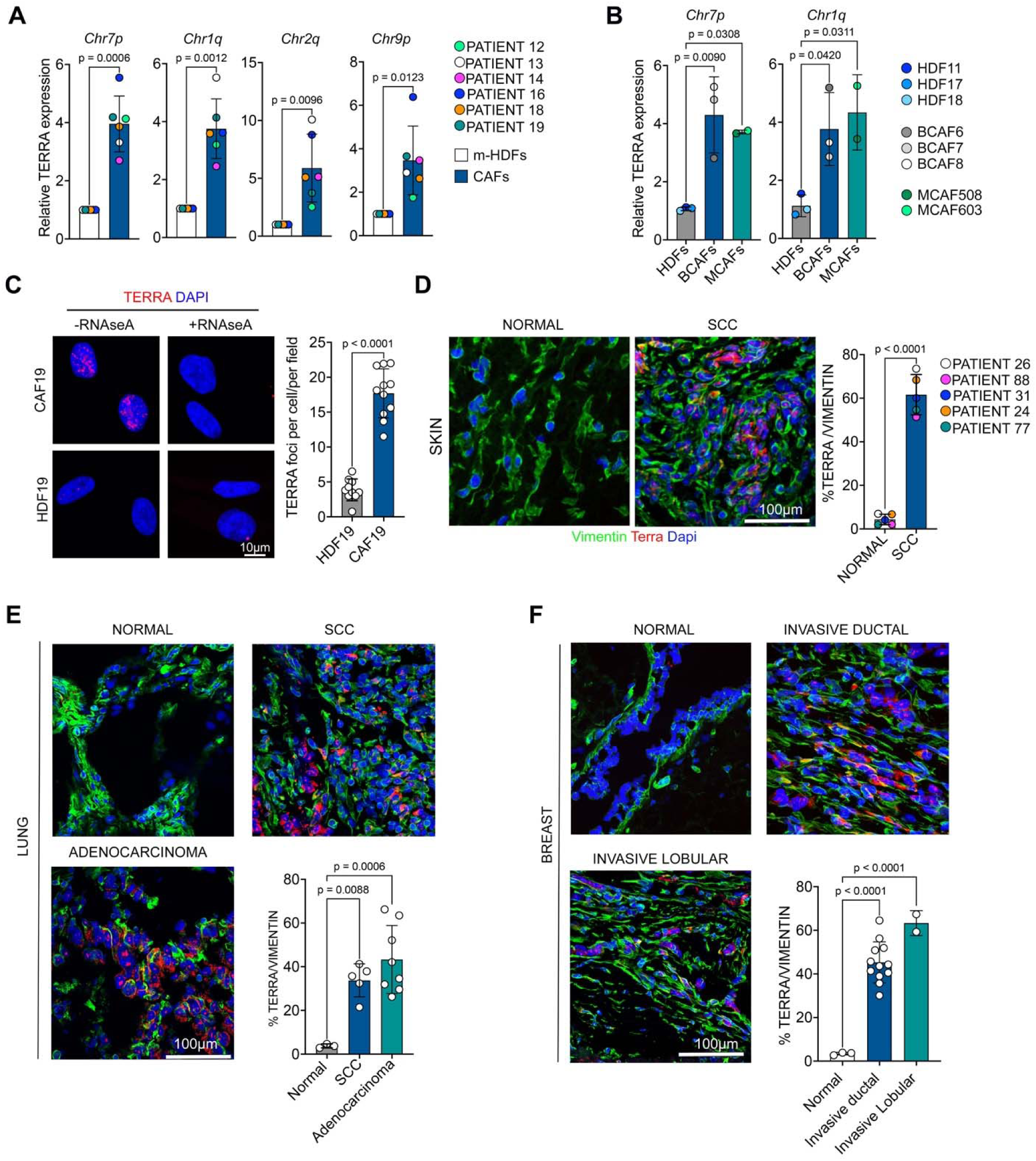
TERRA is highly expressed in CAFs. **(A)** Chromosome-specific TERRA expression in multiple CAF strains isolated from skin SCC lesions (CAFs) compared to HDFs from adjacent unaffected skin (HDFs) from the same patients (#12, #13, #14, #16, #18, #19), as determined by RT-qPCR using specific primers. Expression is shown as fold change in CAFs versus matched HDFs, normalized to GAPDH. n = 6 (strains). Mean ± SD; two-tailed paired t-test. **(B)** RT-qPCR analysis of chromosome-specific TERRA expression in CAFs isolated from BCC lesions (BCAF6, BCAF7, BCAF8) and melanomas (MCAF508, MCAF603) compared to three age-matched reference HDFs (HDF11, HDF17, HDF18) normalized to GAPDH. n = 3 strains (BCAFs) n=2 strains (MCAFs). Mean ± SD. One-way ANOVA with Dunnett’s multiple comparisons test. **(C)** Representative images of TERRA FISH (red) and quantification of TERRA nuclear foci per cell, averaged per field, in CAFs compared to matched HDFs from the same patients. Nuclei were counterstained with DAPI (blue). RNase A treatment (100 µg/ml) was used as a negative control to assess probe specificity. n = 8 to 11 fields per sample. Mean ± SD. Unpaired t-test with Welch’s correction. Quantification of an additional CAF strain is shown in S1D. **(D)** Representative images and quantification of combined immunofluorescence (IF) and RNA FISH analysis of multiple SCC lesions compared to matched adjacent skin from the same patients. IF and RNA FISH were performed using a TERRA probe (red), anti-vimentin antibody (green), and DAPI (blue) for nuclear staining. Quantification represents the percentage of vimentin-positive cells with TERRA foci, averaged from four fields per patient. Images were acquired using a SoRA microscope. n = 5 patients. Mean ± SD. Paired t-test. **(E)** Representative images and quantification of combined immunofluorescence (IF) for vimentin, marking stromal cells, and RNA FISH using a TERRA probe in lung squamous cell carcinoma (SCC), lung adenocarcinoma, and normal lung tissues from a tissue microarray. Full-section vimentin staining is shown in S1F. Quantification indicates the percentage of vimentin-positive cells exhibiting TERRA foci per section. n = 3 normal tissues, n = 5 lung SCC, n = 8 lung adenocarcinoma. Mean ± SD. One-way ANOVA with Dunnett’s multiple comparisons test. **(F)** Representative images and quantification of combined immunofluorescence (IF) for vimentin, marking stromal cells, and RNA FISH using a TERRA probe in breast invasive ductal carcinoma, invasive lobular carcinoma, and normal breast tissues from a tissue microarray. Full-section vimentin staining is shown in S1G. Quantification indicates the percentage of vimentin-positive cells exhibiting TERRA foci per section. n = 3 normal tissues, n = 12 invasive ductal carcinoma, n = 2 invasive lobular carcinoma. Mean ± SD. One-way ANOVA with Dunnett’s multiple comparisons test.

### TERRA is negatively regulated by the androgen receptor (AR)

Downregulation of AR expression provides a well-characterized model for the early steps of CAF activation (*13–15*). One key mechanism regulating TERRA expression is RNA Polymerase II-dependent transcription, which initiates in subtelomeric regions and extends into telomeric repeats (*16*, *17*). Analysis of the nucleotide sequences in several subtelomeric regions revealed the presence of multiple canonical AR binding sites nearby TERRA transcription start sites (Fig. 2A, top; Fig. S2A). Chromatin immunoprecipitation (ChIP) followed by qPCR confirmed AR binding at most of these regions in two independent HDF strains (Fig. 2A, bottom; Fig. S2A, B), suggesting that AR functions as a transcriptional repressor of TERRA. To determine the role of AR in the control of TERRA transcription, we silenced AR in three HDF strains (fig. S3A), resulting in a significant increase in TERRA expression from multiple chromosomes (Fig. 2B; fig. S3B), with RNA-FISH showing increased nuclear TERRA foci that phenocopied the CAF profile (Fig. 2C). To further validate the link between AR activity and TERRA regulation, we used pharmacologic strategies to inhibit AR function. Treatment with ARCC4, a PROTAC compound that induces AR protein degradation (*18*) (Fig. 2D), led to elevated TERRA expression (Fig. 2E, Fig. S3C), with similar effects being elicited by treatment with UT-155, a selective AR antagonist that suppresses AR transcriptional activity (*19*), inducing CAF effector genes (Fig. 2F, Fig. S3D, E, F). Conversely, treatment of CAFs with Ostarine, a selective AR agonist (*20*) that can enhance AR levels via a positive feedback mechanism, reducing CAF activation (*15*) (Fig. 2G), resulted in marked suppression of TERRA expression (Fig. 2H, Fig. S3G). Suppression of TERRA expression was also observed after ostarine treatment of HDFs with shRNA-mediated AR gene silencing (Fig. 2I), which resulted also in an upregulation of endogenous AR expression (Fig. 2J; Fig. S3H). These findings establish AR as a negative regulator of TERRA transcription and demonstrate that AR loss, characteristic of CAFs, directly contributes to TERRA upregulation and fibroblast reprogramming.

**Figure 2.**
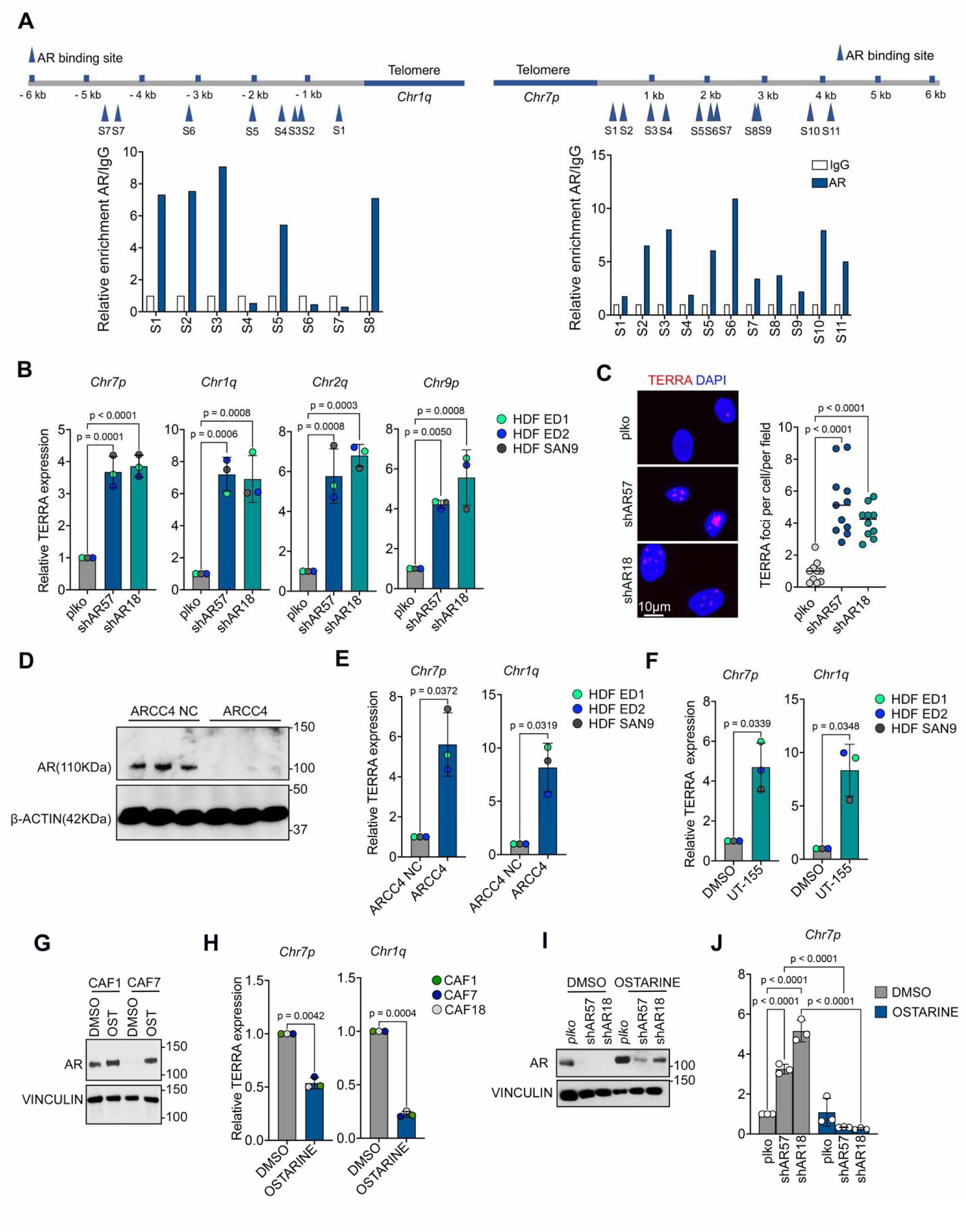
AR is a negative regulator of TERRA transcription. **(A)** Top: Map of predicted AR binding sites (blue arrowheads) within the sub-telomeric 6 kb regions of the indicated chromosomes. Bottom: ChIP-qPCR analysis of AR binding at these sites in HDF1 using AR antibodies or nonimmune IgG controls. Data are expressed as fold enrichment over IgG controls. An independent experiment with a second HDF strain is shown in Supplementary Figure 2b. **(B)** Chromosome-specific TERRA expression in multiple HDF strains following AR silencing by infection with two different shRNAs silencing lentiviruses (shAR#57, shAR#18) versus control (plko), measured by RT-qPCR with specific primers. Expression is shown as fold change relative to control, normalized to GAPDH. n = 3 HDF strains. Data represent mean ± SD. One-way ANOVA with Dunnett’s multiple comparisons test. **(C)** Representative images and quantification of TERRA nuclear foci as assessed by FISH analysis (red) of HDFs with AR silencing (shAR#57, shAR#18) versus control (plko), with DAPI nuclear staining (blue). Data represent pooled fields from 3 independent experiments (n >10 fields total). mean ± SD. One-way ANOVA with Dunnett’s multiple comparisons test. **(D)** Immunoblot analysis with anti-AR and ß-actin antibodies of three HDF strains in charcoal-stripped medium treated with ARCC4 (1 µM, 48 h) versus ARCC4-negative control (ARCC4 NC). **(E)** Chromosome-specific (Chr7p and Chr1q) TERRA expression (in three HDF strains treated with ARCC4 (1 µM, 48 h) versus ARCC4-negative control (ARCC4 NC) as in the previous panel. Expression is shown as fold change relative to ARCC4 NC, normalized to GAPDH. n = 3 strains. Mean ± SD. Paired two-tailed t-test. **(F)** Chromosome-specific (Chr7p, Chr1q) TERRA expression in three HDF strains treated with UT-155 (1 µM, 48 h) versus DMSO control, expressed as fold change relative to DMSO, normalized to GAPDH. n = 3 strains. Mean ± SD. Paired two-tailed t-test. **(G)** Immunoblot analysis of anti-AR and Vinculin antibodies of two CAF strains treated with Ostarine (10 µM, 48 h) or DMSO. **(H)** Chromosome-specific TERRA expression in multiple CAF strains treated with Ostarine (10 µM, 48 h) versus DMSO as in the previous panel. Results are expressed as fold changes relative to DMSO, normalized to GAPDH. n = 3 strains. Mean ± SD. Two-tailed paired t-test. **(I)** Immunoblot analysis with anti-AR and -Vinculin antibodies of HDFs after AR silencing (shAR#57, shAR#18) versus control (plko) treated with Ostarine (10 µM, 48 h) or DMSO. **(J)** RT-qPCR analysis of TERRA expression (Chr 7p) in HDFs after AR silencing (shAR#57, shAR#18) versus control (plko) treated with ostarine (10 µM, 48 h) or DMSO as in the previous panel. Data are expressed as fold change relative to plko with DMSO. Mean ± SD. n = 3 biological replicates. One-way ANOVA with Šídák’s multiple comparisons test.

### Elevated TERRA expression contributes to telomeric damage and transcription control in CAFs

To investigate the functional role of TERRA in the reprogramming of fibroblasts into CAF, we used both loss- and gain-of-function approaches. TERRA was silenced in CAFs using antisense oligonucleotides (TERRA ASO) (*3*) and overexpressed in HDFs using a lentiviral vector (*21*). TERRA knockdown by TERRA ASO transfection led to a clear reduction in TERRA levels across multiple subtelomeric regions compared to scrambled controls (SCR ASO) (Fig. 3A; fig. 4A), along with a significant decrease in nuclear TERRA foci (Fig. 3B). In contrast, overexpression in HDFs increased TERRA foci to levels comparable to those seen in CAFs (Fig. 3C). Notably, neither TERRA silencing nor overexpression affected cellular proliferation (Fig. S4B, C).

**Figure 3.**
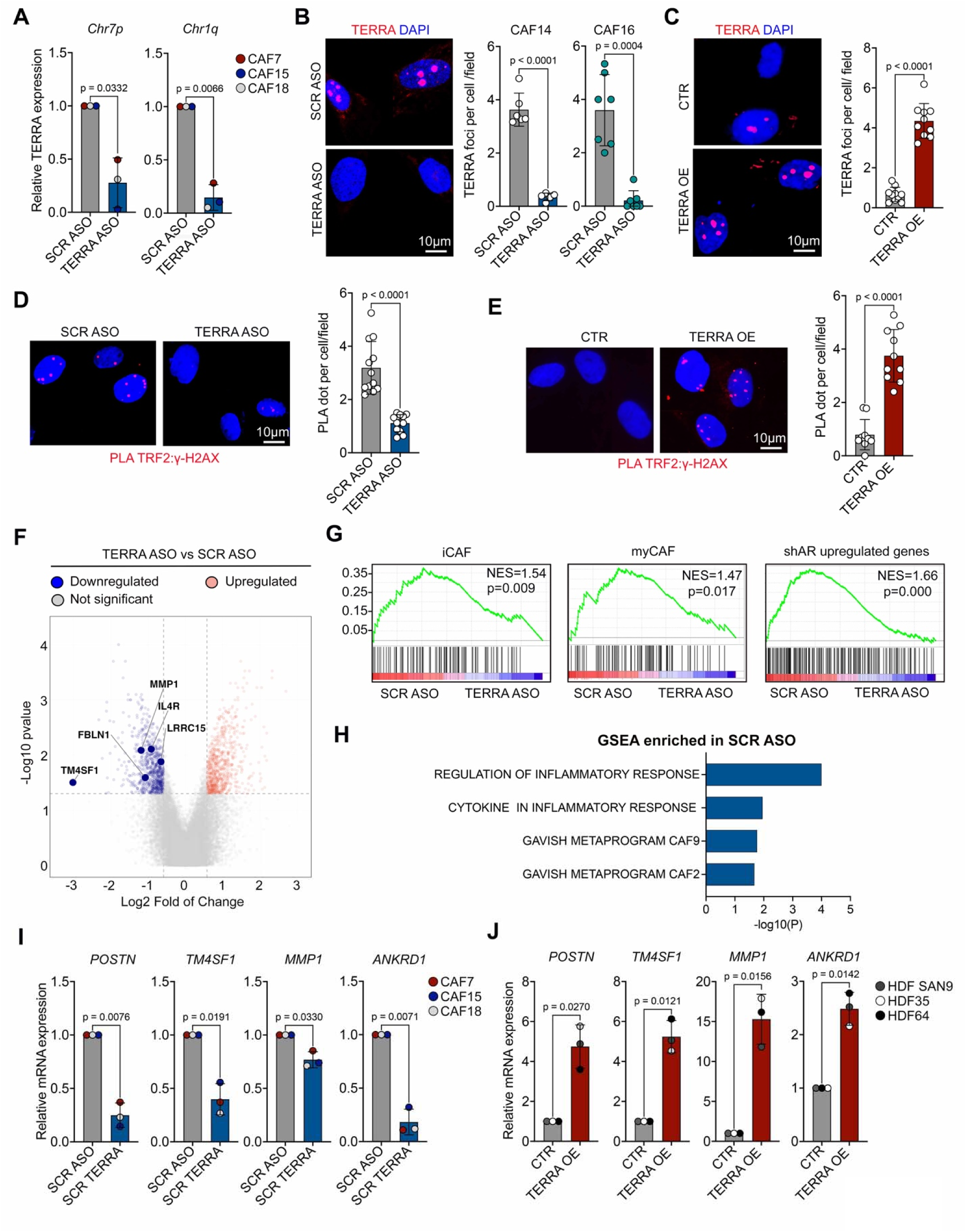
TERRA affects telomeric DNA damage and the CAF transcriptional program. **(A)** Chromosome-specific TERRA expression in CAFs 48 hours after transfection with TERRA ASO or SCR ASO (100 nM). Data are expressed as relative fold change after GAPDH normalization. n=3 strains. Mean ± SD. Two-tailed paired t-test. **(B)** Representative images and quantification of TERRA nuclear foci per cell, averaged per field, detected by RNA FISH (red) in two strains of CAFs transfected with TERRA ASO versus SCR ASO control, as in the previous panel. Data represent pooled fields from 3 independent experiments (n=7 fields total). Mean ± SD. Unpaired t-test with Welch’s correction. **(C)** Representative images and quantification of TERRA nuclear foci per cell, averaged per field, detected by FISH (red) in HDFs infected with a TERRA overexpressing lentiviral vector (TERRA OE) versus a control vector (CTR). Data represent pooled fields from 3 independent experiments (n=10 fields total). Mean ± SD. Unpaired t-test with Welch’s correction. **(D)** Proximity ligation assays (PLA) with antibodies against TRF2 and g-H2AX in CAFs transfected with TERRA ASO or SCR ASO. Shown are representative images and quantification of PLA dots per cell per field. Data represent pooled fields from 3 independent experiments (n=12 fields total). Mean ± SD. Unpaired t-test with Welch’s correction. **(E)** Proximity ligation assays (PLA) with antibodies against TRF2 and γ-H2AX in HDFs upon infection with the TERRA overexpressing lentiviral vector (TERRA OE) versus control vector (CTR). Shown are representative images and quantification of PLA dots per cell per field. Data represent pooled fields from 3 independent experiments (n=10 fields total). Mean ± SD. Unpaired t-test with Welch’s correction. **(F)** Transcriptomic profiling of three CAF strains 48 hours after transfection with TERRA ASO or SCR ASO. Volcano plot displays differential gene expression between TERRA ASO and SCR ASO conditions. The x-axis shows the logO (fold change), and the y-axis shows the –logOO(p-value). Each dot represents a single gene; significantly differentially expressed genes are defined by p < 0.05 and an absolute logO (fold change) ≥ 0.58. Representative downregulated genes with known CAF effector functions are indicated. **(G)** Gene Set Enrichment Analysis (GSEA) of expression profiles from three CAF strains treated with TERRA ASO versus SCR ASO, analyzed against gene signatures associated with inflammatory CAFs (iCAFs), myofibroblastic CAFs (myCAFs), and genes upregulated in HDFs upon AR silencing (*14*). Genes were ranked by signal-to-noise ratio based on their differential expression between SCR ASO and TERRA ASO conditions. The positions of the genes in each set are indicated by black vertical bars, and the enrichment score is shown in green. Normalized enrichment scores (NES) and p-values indicate the degree and significance of enrichment or depletion in TERRA-silenced versus control CAFs. **(H)** Gene Set Enrichment Analysis (GSEA) of the indicated gene signatures from MSigDB. (https://www.gsea-msigdb.org/gsea/msigdb/) including those of the CAF metaprogram CAF9 and CAF2 (*31*), and “Regulation of Inflammatory Response” (GO:0050727) and “Cytokine Production Involved in Inflammatory Response” (GO:0002534). The bar graph shows the - Log10 of p-value in profiles of control versus TERRA-silenced CAF. GSEA with 50 hallmark signatures is shown in Extended Data Fig.4. **(I)** Expression of the indicated CAF effector genes in CAFs transfected with TERRA ASO versus SCR ASO assessed by RT-qPCR. Data are expressed as fold change relative to SCR ASO and normalized to GAPDH. n=3 CAF strains. Mean ± SD. Two-tailed paired t-test. **(J)** Expression of the indicated CAF effector genes in HDFs with TERRA overexpression compared to control, analyzed by RT-qPCR. Data expressed as fold change relative to CTR. Data normalized to RPLP0. n=3 HDF strains. Mean ± SD. Two-tailed paired t-test.

**Fig.4.**
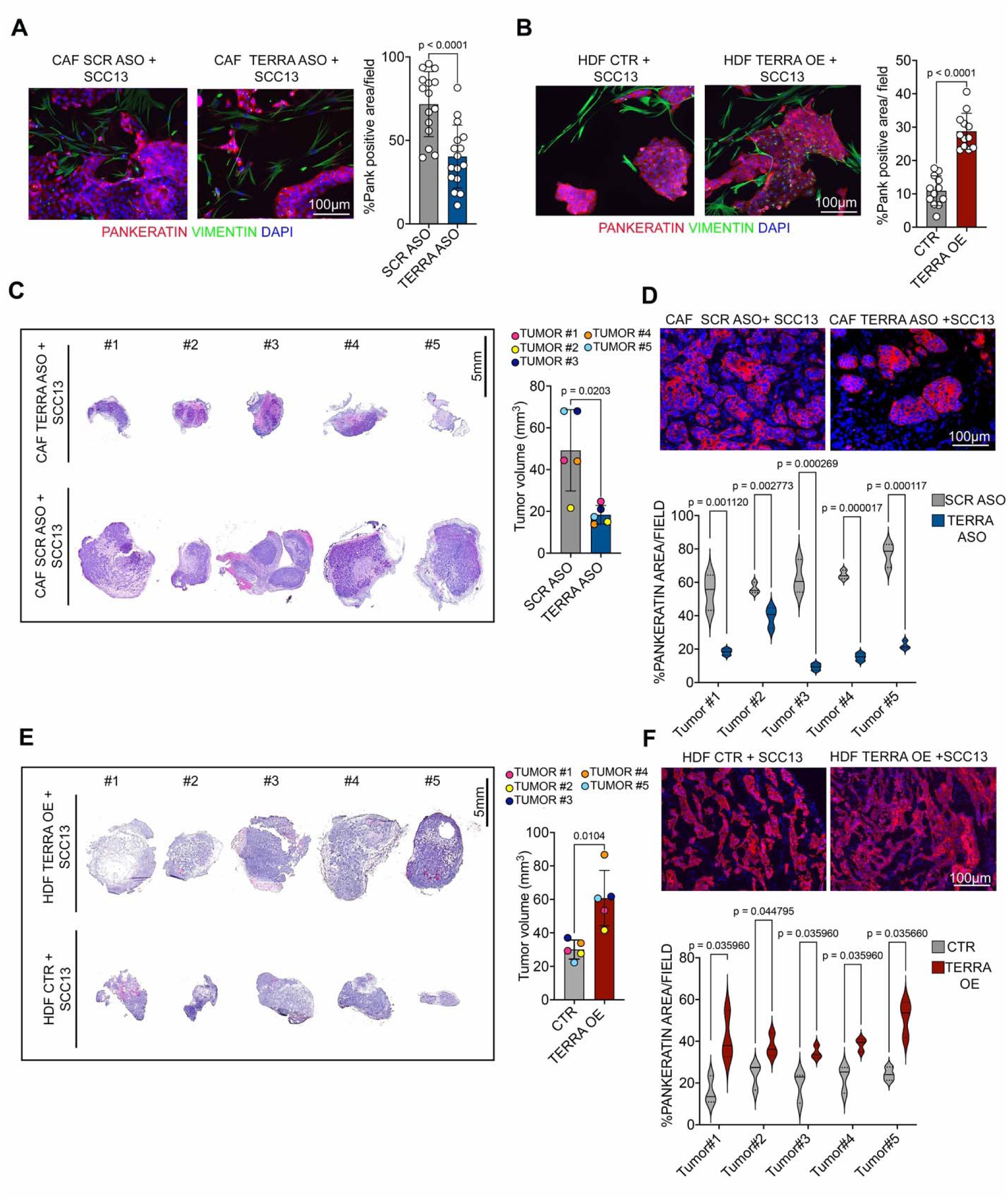
TERRA is required for the proliferation- and tumor-enhancing properties of CAFs on neighboring SCC cells. **(A)** Expansion assays of SCC cells (SCC13) cocultured in a thin Matrigel layer with CAFs transfected with TERRA ASO versus SCR ASO controls (100nM). Shown are representative images of IF analysis with anti-pan-keratin (red) and vimentin (green) antibodies for SCC cells and CAFs identification, respectively, together with quantification of SCC cells expansion, measured by percentage of pan-keratin positive areas. Data represent pooled fields from 3 independent experiments (n=16 fields total). Mean ± SD. Unpaired t-test with Welch’s correction. **(B)** Expansion assays of SCC cells (SCC13) cocultured with HDFs infected with a TERRA overexpressing lentivirus (TERRA OE) versus control virus (CTR), utilizing the same conditions as in the previous panel. Shown are representative images and quantification of the results. Data represent pooled fields from 3 independent experiments (n=12 fields total). Mean ± SD. Unpaired t-test with Welch’s correction. **(C)** Intradermal tumorigenicity assays of SCC13 cells co-injected with CAFs transfected with either TERRA ASO or SCR ASO in contralateral mouse back skin. Shown are representative images of H&E-stained tumor sections along with quantification of tumor volume (mm³), calculated using the formula *(Length × Width²) / 2*. *n =* 5 tumor pairs. mean ± SD. Two-tailed paired t-test. **(D)** Cancer cell density in lesions formed by SCC13 cells admixed with CAFs transfected with either TERRA ASO or SCR ASO, as in the previous panel. Shown are representative images of IF analysis with anti-pan keratin antibodies (red) and quantification of pan-keratin positive area, per lesion. n = 5 tumor pairs, 3 fields per sample. Mean±SD. Multiple unpaired t-test. **(E)** Intradermal tumorigenicity assays of SCC13 cells co-injected with HDFs infected with either a TERRA overexpression (TERRA OE) or control (CTR) in the contralateral mouse back skin. Shown are representative images of H&E-stained tumor sections along with quantification of tumor volume (mm³), calculated using the formula *(Length × Width²) / 2*. *n =* 5 tumor pairs. mean ± SD. Two-tailed paired t-test. **(F)** Cancer cell density in lesions formed by SCC13 cells admixed with TERRA overexpressing HDFs versus controls as in the previous panel. Shown are representative images of IF analysis with anti-pan keratin antibodies (red) and quantification of pan-KRT positive area, per lesion. n = 5 tumor pairs, 3 fields per sample. Mean±SD. Multiple unpaired t-test.

We previously showed that CAFs exhibit persistent telomeric damage (*22*, *23*). To determine whether TERRA contributes to this phenotype, we performed proximity ligation assays (PLAs) using antibodies against γ-H2AX, a marker of DNA damage, and TRF2, a telomeric protein, to specifically detect telomeric breaks (*24*). Consistent with previous findings in which the suppression of TERRA transcription alleviates the telomere instability, we found that silencing TERRA in CAFs significantly reduced telomeric DNA damage. (Fig. 3D). Conversely, TERRA overexpression in HDFs was sufficient to induce telomeric damage (Fig. 3E).

Beyond its impact on telomere integrity, we investigated whether elevated TERRA expression contributes to the transcriptional reprogramming of cancer-associated fibroblasts (CAFs). Global transcriptomic analysis revealed a core set of genes consistently up- or downregulated across three independent CAF strains following TERRA silencing (Fig. 3F). Several established CAF effector genes, including *LRRC15* (*25*)*, MMP1* (*26*)*, IL4R* (*27*)*, and FLN1* (*28*), were significantly downregulated upon TERRA depletion. TM4SF1, coding for a recently identified CAF effector involved in both skin (*29*) and gastric (*30*) cancer, was among the most strongly suppressed.

Gene Set Enrichment Analysis (GSEA) further revealed that TERRA knockdown led to coordinated downregulation of multiple CAF-related gene signatures, including inflammatory and myofibroblastic CAF subtypes, a transcriptional program associated with AR loss (*14*), two CAF metaprograms from an extensive analysis of intratumor heterogeneity across multiple cancer types (*31*), and gene sets linked to inflammation and cytokine signaling (Fig. 3G, H). A set of other functionally relevant gene signatures were also affected by TERRA silencing (fig. S4d). Direct RT–qPCR analysis confirmed that TERRA silencing reduced expression of key CAF effectors, including *POSTN, MMP1, TM4SF1,* as well as *ANKRD1*, coding for a transcriptional co-activator implicated in CAF activation (*14*) (Fig. 3i). Conversely, ectopic TERRA overexpression in HDFs induced upregulation of these same genes (Fig. 3J).

Together, these findings provide functional evidence that elevated TERRA promotes both telomeric instability and transcriptional reprogramming in CAFs, thereby actively contributing to CAF activation and tumor–stroma remodeling.

### TERRA promotes the pro-tumorigenic activity of CAFs

A hallmark of CAFs is their ability to promote the proliferation and tumorigenicity of adjacent cancer cells. To investigate the functional significance of TERRA in this context, we initially utilized a cancer–stromal cell co-culture assay embedded in a thin Matrigel layer (*13*). The expansion of squamous carcinoma cells (SCC13) was markedly reduced in the presence of TERRA-silenced CAFs relative to controls (Fig. 4A), whereas it was enhanced in the presence of TERRA-overexpressing HDFs (Fig. 4B).

In an orthotopic cancer model involving intradermal injection of cells into the back skin of mice, tumors formed by SCC cells co-injected with TERRA-silenced CAFs were significantly smaller and exhibited lower cancer cell density compared to those formed with control CAFs (Fig. 4C, D). Conversely, SCC cells co-injected with TERRA-overexpressing HDFs gave rise to larger, more cellular tumors than those admixed with control HDFs (Fig. 4E, F). These findings demonstrate that elevated TERRA expression in CAFs promotes cancer cell proliferation and tumor growth.

### NONO Differentially Associates with TERRA and AR in CAFs

To identify TERRA-interacting proteins that may contribute to CAF activation, we performed iDRIP (identification of direct RNA-interacting proteins) (*3*) in CAFs using with an antisense probe to pull down TERRA and a sense probe as a negative control (Fig. 5A). Quantitative mass spectrometry across two biological replicates identified 19 proteins significantly enriched in TERRA pulldowns (Table S1), predominantly RNA-binding proteins involved in transcription, splicing, and ribosome biology, and others related to cytoskeletal organization (fig.S5A). Among the enriched proteins, we focused on NONO, a multifunctional RNA- and DNA-binding protein previously shown to interact with TERRA and regulate its localization at telomeres (*6*). STRING network analysis (https://string-db.org/) revealed high-confidence interactions between NONO and several other RNA-binding proteins identified in the pulldown (Fig. 5b), consistent with its role as a molecular scaffold involved in transcription, RNA processing, and genome stability (*5*). RNA immunoprecipitation (RIP) using NONO-specific antibodies confirmed its binding to TERRA transcripts across multiple CAF strains (Fig. 5C, fig. S5B). Immuno-RNA FISH with anti-NONO antibodies and a TERRA probe, combined with super-resolution (SoRa) microscopy, revealed frequent nuclear colocalization between NONO and TERRA in CAFs, with over 50% of foci overlapping (fig. S5C).

**Fig.5.**
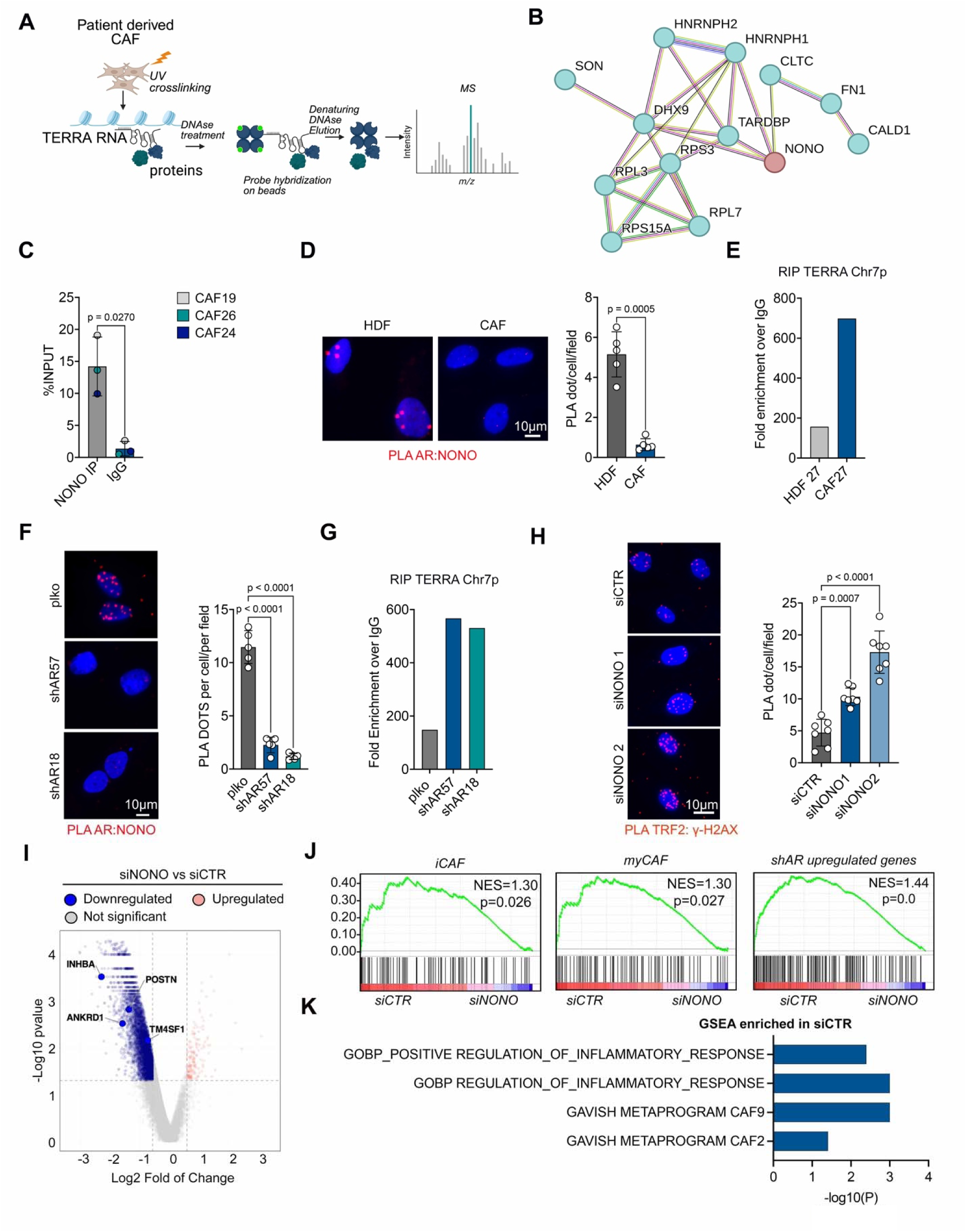
NONO associates with TERRA or AR in CAFs versus HDFs. **(A)** Schematic representation of the iDRIP-MS (identification of direct RNA-interacting proteins – Mass Spec) workflow used to identify TERRA-interacting proteins in CAFs (Biorender). **(B)** Predicted interacting network of the identified TERRA-associated proteins based on experimental evidence, curated databases, or co-expression by STRING (https://www.expasy.org/resources/string), using a confidence score threshold of 0.4. The type of connections is denoted by lines of different colors. Known interactions (light blue, magenta), predicted interactions (green, red, blue), others (black, purple, lime). Note the NONO protein (in red) as one center of the network. **(C)** TERRA-NONO association in multiple CAF strains (CAF#19, CAF#24, CAF#26), as assessed by RNA immunoprecipitation (RIP) with antibodies against NONO versus non-immune IgGs, followed by RT-qPCR with primers specific to TERRA transcripts from chromosome 7p. Data are expressed as percentage of input. Mean±SD, paired two-tailed *t*-test. **(D)** PLA assay detecting endogenous AR/NONO interactions in CAFs versus matched HDFs. Red puncta indicate AR/NONO complexes; nuclei were counterstained with DAPI (blue). Quantification represents the number of PLA dots per cell per field. n = 5 fields per sample (counting > 20 cells per field), shown as mean ± SD. Unpaired t-test with Welch’s correction. **(E)** RIP assay performed using an antibody against NONO or control IgG in CAF or matched HDF (#27). TERRA levels from chromosome 7p were quantified by RT-qPCR and normalized to input. Data are presented as fold enrichment over control IgG. **(F)** NONO-AR association as detected by Proximity ligation assays (PLAs) of HDFs of HDFs plus/minus shRNA-mediated AR silencing as a control of specificity. Shown are representative images of nuclear NONO-AR complexes (red dots) together with quantification of PLA dots per cell per field. n = 5 fields (counting > 20 cells per field). Mean ± SD, one-way ANOVA with Dunnett’s multiple comparisons test. **(G)** RNA immunoprecipitation (RIP) was performed using an antibody against NONO or control IgG in HDFs transduced with control lentivirus (plko) or AR-targeting shRNAs (shAR57 and shAR18). TERRA levels from chromosome 7p were quantified by RT-qPCR and normalized to input. Data are presented as fold enrichment over control IgG. **(H)** PLA assays with antibodies against TRF2 and γ-H2AX in CAFs at 72 hours after transfection with NONO targeting siRNAs versus controls as in the previous panel. Shown are representative images and quantification of the number of PLA puncta per cell per field. n =7 fields per sample (counting > 20 cells per field). Mean ± SD, one-way ANOVA with Dunnett’s multiple comparison correction. **(I)** Transcriptomic profiling of two CAF strains 72 hours after transfection with NONO targeting siRNAs versus controls. Shown is a volcano plot of differentially expressed genes with logOfold change (x-axis) and–logOOp-value (y-axis). Each dot represents a single gene; significantly differentially expressed genes are defined by p < 0.05 and an absolute logO (fold change) ≥ 0.58. Representative downregulated genes with known CAF effector functions are indicated. **(J)** Gene Set Enrichment Analysis (GSEA) plot using expression profiles from two CAF strains treated with siCTR or siNONO, analyzed against gene sets associated with inflammatory CAFs (iCAFs), myofibroblastic CAFs (myCAFs), and genes upregulated in HDFs upon AR silencing (*14*). Genes were ranked by signal-to-noise ratio based on their differential expression between siCTR and siNONO conditions. The positions of the genes in each set are indicated by black vertical bars, and the enrichment score is shown in green. Normalized enrichment scores (NES) and p-values indicate the degree and significance of enrichment or depletion in NONO-silenced CAFs.

NONO has been previously reported to associate with AR in prostate cancer cells to regulate gene transcription (*32*, *33*). Given that AR acts as a negative regulator of both TERRA expression and CAF activation, we investigated whether it also modulates the formation of the TERRA–NONO complex. While AR expression is reduced in CAFs relative to matched HDFs (*13*), NONO protein levels remained constant (Fig. S5D). Proximity ligation assays (PLA) revealed abundant nuclear NONO–AR complexes in HDFs, which were markedly diminished in CAFs (Fig. 5D), coinciding with a reciprocal increase in NONO–TERRA interactions (Fig. 5E). To directly test the role of AR, we evaluated the effects of its silencing in HDFs. This led to a loss of NONO–AR complexes (Fig. 5F) without affecting NONO protein abundance (fig. S5E), but significantly enhanced NONO-TERRA association (Fig. 5G). Together, these findings reveal a dynamic and context-dependent interaction between NONO, AR, and TERRA that shifts during CAF activation. A model can be proposed in which AR sequesters NONO in HDFs and limits its interaction with TERRA, whereas loss of AR in CAFs releases NONO to form complexes with TERRA, facilitating the transcriptional reprogramming characteristic of the CAF state.

### NONO is a determinant of the transcriptional program of CAF activation

To investigate the functional role of NONO in CAFs, we performed siRNA-mediated silencing, achieving efficient knockdown by 72 hours (fig. S5F). This led to a marked increase in telomeric DNA damage (Fig. 5H), consistent with the reported role of NONO in maintaining telomere integrity in antagonism with TERRA (*6*). To determine whether NONO also contributes to gene regulation in CAFs, we conducted transcriptomic profiling of multiple CAF strains following NONO depletion using two independent siRNAs. Knockdown of NONO induced widespread transcriptional alterations, including downregulation of established CAF effector genes (*POSTN*, *INHBA*, *TM4SF1*) and transcriptional regulators of CAF identity such as *ANKRD1* (*14*) (Fig. 5I), closely resembling the effects of TERRA silencing (Fig. 3I). These changes were validated by RT–qPCR and immunoblotting (fig. S5G, H).

Gene Set Enrichment Analysis (GSEA) further demonstrated that NONO silencing significantly suppressed transcriptional programs associated with inflammatory and myofibroblastic CAFs, a CAF signature induced by AR loss (*14*), and two CAF metaprograms (Fig. 5J, K), all of which were similarly reduced by TERRA depletion (Fig. 3G, H). In line with its broader functional roles, more global GSEA revealed that NONO knockdown affected a wider array of CAF-related gene signatures than TERRA, suggesting additional roles in cellular processes beyond those regulated by TERRA (fig. S6).

Together, these findings identify NONO as a key regulator in CAFs, essential for sustaining their pro-tumorigenic transcriptional program, with overlapping but broader functions than TERRA.

### Pharmacologic targeting of the TERRA–NONO complex reverses CAF activation, including in human clinical lesions

To assess the translational relevance of our findings, we evaluated the effects of the small-molecule (R)-SKBG-1, which covalently binds to Cys145 in NONO, altering its RNA-binding properties and impairing its transcript-processing function (*32*). Treatment of CAFs with (R)-SKBG-1 downregulated key effector and regulatory genes (*POSTN*, *INHBA*, *TM4SF1*, *ANKRD1*), effectively phenocopying the effects of NONO depletion (Fig.6A). (R)-SKBG-1 also prevented the induction of these genes in HDFs following *AR* silencing, mimicking the effects observed with TERRA knockdown (Fig. 6B, Fig. S7A)

**Figure 6.**
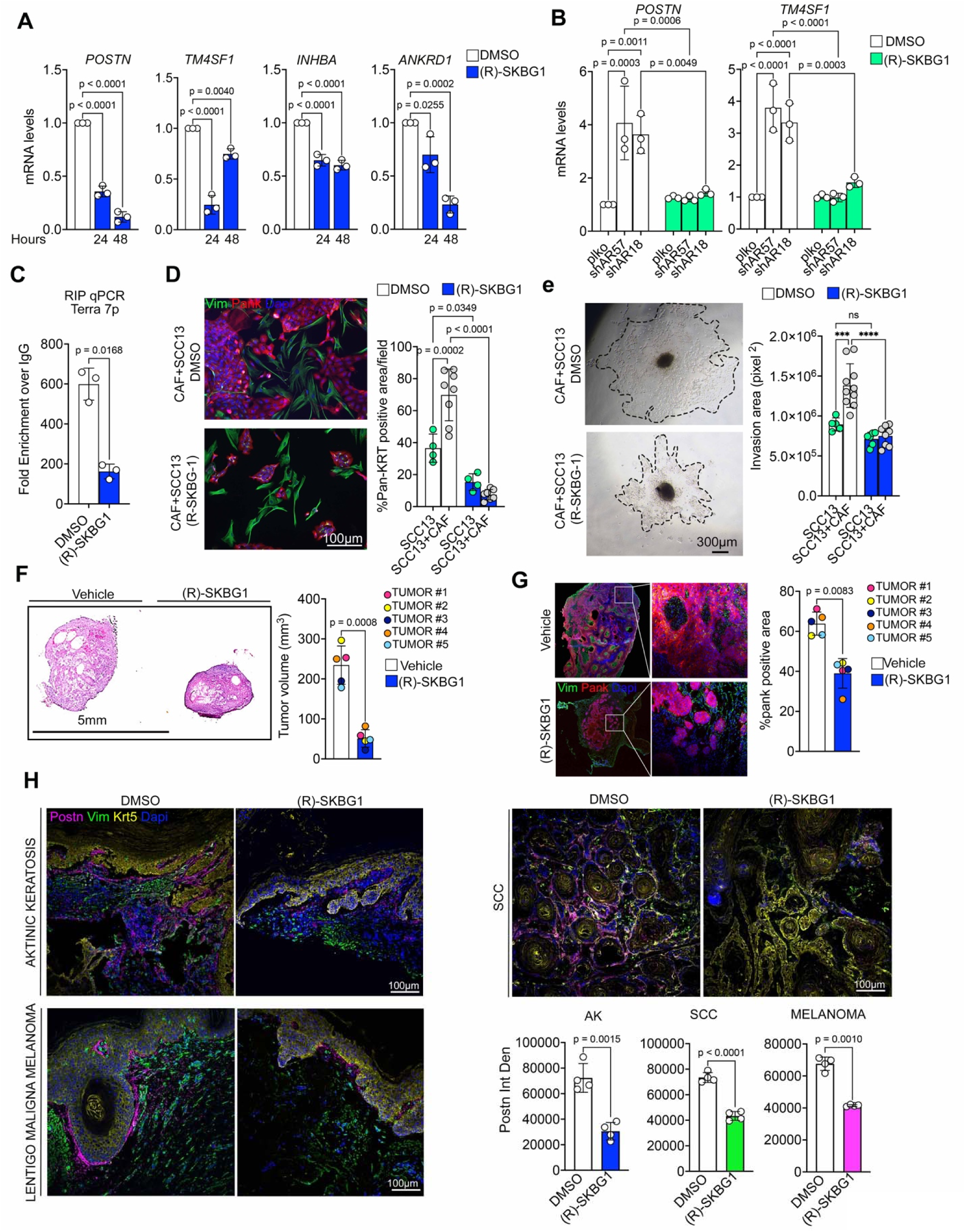
Pharmacological inhibition of NONO disrupts the TERRA–NONO complex and suppresses CAF-driven tumorigenesis. **(A)** Expression of the indicated CAF effector genes in CAFs with DMSO or (R)-SKBG1 treatment (5 μM) after 24, 48 hours, analyzed by RT-qPCR. Data expressed as fold change relative to DMSO. Data normalized to RPLP0.n=3 biological replicates. Mean ± SD. One-way ANOVA with Dunnett’s multiple comparison test. **(B)** Expression of the indicated CAF effector genes in HDFs plus/minus AR silencing, followed by DMSO or (R)-SKBG1 treatment (5mM) for 48 hours analyzed by RT-qPCR. Data expressed as fold change relative to DMSO. Data normalized to RPLP0.n=3 biological replicates. Mean ± SD. One-way ANOVA with Šídák’s multiple comparisons test **(C)** RIP assay performed using an antibody against NONO or control IgG in CAF treated with DMSO or (R)-SKBG1 as described before. TERRA levels from chromosome 7p were quantified by RT-qPCR and normalized to input. Data are presented as fold of enrichment over control IgG. n=3 biological replicates. Mean ± SD. Paired two-tailed t-test. **(D)** Expansion assays of SCC cells (SCC13) alone or cocultured with CAFs treated with (R)-SKBG1 versus DMSO as above. Shown are representative images of immunofluorescence analysis with anti-pan-keratin (red) and anti-vimentin antibodies (green) for SCC13 and CAF identification, respectively, together with quantification of SCC cells expansion, measured by percentage of pan-keratin positive areas. Data represent pooled fields from 3 independent experiments (n>4 fields total). Mean ± SD. One-way ANOVA with Šídák’s multiple comparisons test. Note the significantly higher expansion of SCC cells admixed with CAFs relative to when tested in isolation, and the even greater suppression caused by (R)-SKBG1 treatment in the presence of CAF. **(E)** Spheroid invasion assays of SCC cells alone or admixed with CAFs and treated with DMSO or (R)-SKBG1 as described before. Shown are representative bright field images of spheroids and quantification of the invasion area measured as the difference between the core and the surrounding invaded area delimited with dotted lines. n >5 spheroids per condition. Mean ± SD. One-way ANOVA with Šídák’s multiple comparisons test. ns=0.3126 Note the significantly higher invasion capability of SCC cells admixed with CAFs relative to when tested in isolation, and the suppression by (R)-SKBG1 treatment. **(F)** SCC13 cells were co-injected intradermally with CAFs into the contralateral flanks of immunocompromised mice. Upon detection of palpable lesions (∼5 mm in diameter) 7 days post-injection, (R)-SKBG1 (10 µM diluted in PBS) or vehicle (PBS) was administered intratumorally into paired lesions once daily for 14 days. Shown are representative images of H&E-stained tumor sections and quantification of tumor volume (mm³), calculated using the formula (length × width²) / 2. n = 5 tumor pairs; data represent mean ± SD; statistical analysis was performed using a two-tailed paired t-test. **(G)** Cancer cell density in lesions formed by SCC13 cells admixed with CAFs treated with (R)-SKBG1 or PBS as in the previous panel. Shown are representative images of IF analysis with anti-pan keratin antibodies (red) to visualize SCC13 cells and with anti-vimentin (green) for SCC13 and CAF identification, respectively. Quantification of pan-keratin positive area, per lesion. n = 5 tumor pairs. Mean±SD, paired two-tailed *t*-test. **(H)** Representative immunofluorescence images of actinic keratosis (AK), squamous cell carcinoma (SCC), and lentigo maligna melanoma biopsies treated ex vivo with DMSO or the NONO inhibitor (R)-SKBG1 (10 µM) for 48 h. Sections were stained for periostin (POSTN; magenta), vimentin (Vim; green), keratin 5 (Krt5; yellow), and DAPI (blue). Lower panels: quantification of POSTN immunofluorescence signal intensity (integrated density) in AK, SCC, and melanoma lesions. Each point represents one field. Mean ± SD; Unpaired t test with Welch’s correction.

Importantly, (R)-SKBG-1 treatment did not alter overall NONO or TERRA expression levels (fig. S7B, C) but significantly impaired the ability of NONO to bind TERRA, as determined by RNA immunoprecipitation (RIP) in CAFs (Fig. 6C), pointing to TERRA–NONO complex formation as a primary target.

(R)-SKBG-1 has previously been shown to inhibit proliferation in various cancer cell types (*32*, *34*, *35*). In our assays, (R)-SKBG-1 reduced SCC cell proliferation in monoculture, with more pronounced effects in co-culture with CAFs, where it suppressed CAF-mediated cancer cell expansion (Fig. 6d). In 3D tumor spheroid models, CAFs promoted SCC invasion, which was markedly attenuated by (R)-SKBG-1 treatment (Fig. 6E), phenocopying the effects of NONO silencing using two independent siRNAs (Fig. S7D).

To evaluate the impact of (R)-SKBG-1 *in vivo*, including its ability to reverse established tumors, SCC cells were co-injected intradermally with CAFs into immunodeficient mice. Once palpable tumors had formed, animals received intratumoral injections of (R)-SKBG-1 or vehicle control for one week. As shown in Fig. 6F and Fig. S7E, (R)-SKBG-1 treatment resulted in a marked reduction in tumor volume and a significant decrease in cancer cell density (fig.6G).

To assess the clinical potential of the findings, we treated freshly excised actinic keratosis, cSCC, and melanoma lesions with (R)-SKBG-1 (fig. S7F). This treatment markedly reduced expression of the CAF marker POSTN across all three lesion types (fig.6H), indicating its potential as a potent therapeutic strategy to attenuate CAF activation in patients. Levels of FAP, another established CAF marker, were similarly decreased (fig. S7G), further supporting the translational potential of targeting this pathway to reverse CAF activation in the clinical setting.

Collectively, these findings demonstrate that pharmacologic targeting of the TERRA–NONO complex with (R)-SKBG-1 effectively reprograms CAFs, disrupts their pro-tumorigenic functions, and limits tumor growth in vivo. The observed reduction of CAF markers in freshly excised pre-malignant and malignant human lesions underscores the translational potential of this approach and positions (R)-SKBG-1 as a promising therapeutic candidate for stroma-directed cancer therapy.

## Discussion

We have identified a previously unrecognized telomere-linked mechanism that orchestrates CAF activation and function. The lncRNA TERRA, transcribed from telomeric regions, is up-regulated in CAFs and drives widespread transcriptional reprogramming through its interaction with the multifunctional RNA-binding protein NONO. While elevated TERRA expression contributes to increased telomeric DNA damage in CAFs, NONO exerts a counterbalancing protective role. At the transcriptional level, however, TERRA and NONO converge to promote a tumor-promoting gene expression program. Genetic suppression or pharmacological disruption of the TERRA-NONO interaction reverses the CAF phenotype and reveals a targetable vulnerability to counteract cancer–stroma cells expansion and progression.

TERRA expression is regulated by a complex interplay of transcription factors, epigenetic modifications, and telomere-binding proteins (*16*, *36*, *37*). Here, we identify AR, a known context-dependent transcriptional activator and repressor (*38*), as a direct regulator of TERRA transcription. AR binds to TERRA promoters at multiple chromosome ends, and its loss in HDFs leads to TERRA upregulation. Conversely, reactivation of AR in CAFs, where AR levels are diminished, suppresses TERRA expression. Functionally, AR silencing in HDFs mirrors the effects of TERRA overexpression, inducing telomeric DNA damage (Extended Data Fig. 8) while TERRA silencing in CAFs reduces telomeric stress. Together, these findings support a model in which AR downregulation, such as that induced by UVA exposure (*13*), a driver of skin aging and carcinogenesis (*39*), triggers a telomeric stress response marked by TERRA accumulation, ultimately contributing to fibroblast reprogramming into CAFs.

Importantly, TERRA is not merely a marker of telomere dysfunction in CAFs but an active driver of their transcriptional identity. TERRA depletion reprograms CAFs toward a more HDF-like state and suppresses their tumor-promoting activity, whereas increased TERRA expression in HDFs induces a CAF-like phenotype. A central mechanistic insight from our study is the identification of NONO as a direct TERRA-binding partner essential for this reprogramming. While NONO has been previously implicated in diverse contexts (*5*), it has not been studied in cancer-associated stromal cells. We find that NONO binds TERRA to a much greater extent in CAFs than in HDFs and is required to sustain CAF-specific transcription programs, including effector and regulatory genes. Notably, NONO depletion increases telomeric DNA damage in CAFs, while TERRA silencing mitigates it, consistent with their antagonistic roles at telomeres (*6*). AR acts as a key upstream modulator: in HDFs, it forms nuclear complexes with NONO, restricting its availability to engage with TERRA. AR loss, a hallmark of CAF activation (*13–15*), frees NONO to form TERRA–NONO complexes, thereby promoting the transcriptional and functional reprogramming into CAFs.

The translational relevance of these findings is underscored by our demonstration that a small-molecule NONO inhibitor, (R)-SKBG-1 (*32*), disrupts the NONO-TERRA interaction and recapitulates the effects of NONO depletion. (R)-SKBG-1 suppresses CAF effector gene expression and impairs stromal-enhanced cancer cell growth in 3D co-culture and spheroid invasion models. (R)-SKBG-1 has been previously shown to suppress NONO function by binding to this protein, covalently modifying it at a specific residue in a hinge region between two RNA recognition motifs (RRMs) and proximal to residues critical for RNA binding and affecting its RNA binding properties (*32*). In the context of CAF activation, we found that (R)-SKBG-1 did not affect TERRA expression or overall NONO protein levels but disrupted TERRA–NONO association and complex formation, consistent with a specific mode of action targeting RNA–protein interactions rather than gene expression or protein stability. Importantly, treatment of freshly excised human biopsies of actinic keratosis, cSCC, and melanoma with (R)-SKBG-1 markedly reduced the expression of the CAF markers POSTN and FAP, providing direct evidence that targeting the TERRA–NONO complex can reverse CAF activation in patient-derived tissues and highlighting the therapeutic promise of this approach.

Together, our findings reveal a previously unrecognized telomere-derived RNA–protein axis that governs fibroblast identity and function through the interplay of TERRA, NONO, and AR. More broadly, they highlight the capacity of telomere-associated noncoding RNAs to act as dynamic regulators of the tissue microenvironment, with potential relevance not only to cancer but also to fibrosis, chronic inflammation, and aging-associated tissue degeneration. By linking telomere dysfunction to stromal remodeling through specific RNA–protein interactions, our work opens new conceptual and therapeutic avenues for targeting fibroblast-driven pathologies.

## Methods

### Human Subjects

Primary cancer-associated fibroblasts (CAFs) and matched normal fibroblasts (NFs) were isolated from discarded skin specimens of cutaneous squamous cell carcinoma (SCC) patients undergoing surgery at the Department of Dermatology, Massachusetts General Hospital (Boston, MA, USA), under institutional approval (Protocol #2018P003156).

Shave biopsies of actinic keratosis and lentigo malignant melanoma were provided by the Dermatology Department of the University Hospital Zürich (Switzerland) with the assistance of the SKINTEGRITY.CH biobank. The use of human samples for research was approved by the local ethics commission (BASEC_Nr.=2017-00688). All samples used were surplus materials from routine surgeries.

### MouseModels

NOD.CB17-Prkdc^scid^ Il2rg^tm1Wjl^/SzJ (NSG) mice were obtained from The Jackson Laboratory (Stock No. 005557) and maintained under specific pathogen-free (SPF) conditions in the Massachusetts General Hospital (MGH) animal facility. Both male and female mice, 6–10 weeks of age at the start of experiments, were used. Mice were housed in individually ventilated cages with a 12-hour light/dark cycle, controlled temperature (20–22°C) and humidity (40–60%), and provided ad libitum access to autoclaved food and water. All animal experiments were conducted in compliance with the guidelines of the MGH Institutional Animal Care and Use Committee (IACUC) and approved under protocol (2004N000170).

### Isolation of Cancer-Associated Fibroblasts from Skin

Cancer-associated fibroblasts (CAFs) were isolated from discarded skin specimens of squamous cell carcinoma (SCC), alongside matched normal fibroblasts (NFs) derived from adjacent non-tumoral skin at the Department of Dermatology, Massachusetts General Hospital (Boston, Massachusetts, USA) with institutional approval (2018P003156). Briefly, after surgical removal of adipose tissue, the biopsies were minced into 1–2 mm fragments and digested in 0.25 mg/mL Liberase TL (Roche, Cat# 05401020001) at 37°C for 40 minutes. The enzymatic reaction was quenched with fetal bovine serum (FBS), and the cell suspension was filtered through a 70Oµm strainer using a syringe. Cells were pelleted by centrifugation at 300 × g and plated in complete DMEM containing 10% FBS, 1% penicillin, and 1% streptomycin. Cultures were maintained by splitting at 80–90% confluence, with medium changes every 48 hours. Multiple vials were cryopreserved at passage 2, and all experiments were performed using early-passage CAFs (passages 3–6). Cell strains were routinely tested for Mycoplasma contamination. CAFs from basal cell carcinoma (BCAFs) and melanomas (MCAFs) were previously obtained (*13*).

### Isolation of Human Dermal Fibroblasts (HDFs)

Human dermal fibroblasts (HDFs) were isolated from discarded foreskin or abdominoplasty skin samples obtained from the Department of Dermatology at Massachusetts General Hospital (Boston, Massachusetts, USA) under institutional approval (protocol #2021P001670) or were previously established as described (*40*). Multiple vials were cryopreserved at passage 2, and all experiments were performed using early-passage HDFs (passages 3–6).

### Cutaneous SCC cell lines

SCC cells were extracted from the skin and cultured as previously described (*41*). SCC13 cells were kindly provided by Dr. James Rheinwald (Brigham and Women’s Hospital, Boston, MA, USA). Cells were cultured in DMEM supplemented with 10% fetal bovine serum (FBS) and 1% penicillin-streptomycin.

### Lentiviral Production and Transduction

Lentiviral particle production and infections were performed as previously described (*42*). Briefly, HEK 293T cells were seeded to reach ∼60% confluence. After 24 hours, cells were transfected using polyethylenimine (PEI) and the appropriate vector in DMEM supplemented with 10% fetal bovine serum (FBS), without antibiotics. For transfection of a 15 cm dish, 14 µg of total DNA (comprising the vector of interest and packaging plasmids—CMG and VSV-G—in a 4:2:1 ratio) were mixed with 42 µL of PEI (maintaining a 1:3 DNA:PEI ratio), vortexed, and incubated for 15 minutes at room temperature. During this incubation, the culture medium was replaced with fresh DMEM containing 10% FBS (without antibiotics). The transfection mix was then added to the cells and incubated for 12 hours, after which the medium was replaced with complete DMEM. Viral supernatants were collected 48 hours post-transfection, filtered through 0.45 µm filters, and stored in aliquots at –80°C. Target cells were transduced by overnight incubation with the viral supernatant. The following morning, the medium was replaced with complete DMEM. After 48 hours, antibiotic selection was initiated to enrich for transduced cells.

### shRNA-Mediated AR Silencing

CAFs and HDFs were transduced with lentiviral shRNA vectors developed by The RNAi Consortium (TRC). HDFs were infected with two independent shRNA constructs targeting the androgen receptor (AR): shAR#1 (TRCN0000003718) and shAR#2 (TRCN0000003715), or with the empty pLKO vector as a control. All constructs included a puromycin resistance gene for antibiotic selection. Vector sequences are listed in Supplementary Table 1.

### TERRA Overexpression

Human dermal fibroblasts (HDFs) were transduced with a lentiviral vector encoding approximately 130 telomeric hexanucleotide repeats consisting of TTAGGG and TTGGGG variant sequences (pLV CMV-TERRA; Addgene plasmid #111375, RRID: Addgene_111375; http://n2t.net/addgene:111375; doi:10.3390/genes7080046). A control vector, pLenti CMV/TO Zeo DEST 644-1 (Addgene plasmid #17249), was used as a negative control. Both vectors contain a zeocin resistance gene for antibiotic selection. Vector sequences are listed in Table S2.

### siRNA-Mediated NONO silencing

Cancer-associated fibroblasts (CAFs) were transfected with two independent siRNAs targeting NONO(siNONO#1(Mission siRNA NM_007363 SASI_Hs02_00343477 and siNONO#2 (Mission siRNA NM_007363 SASI_Hs02_00343479 or with a non-targeting control siRNA (MISSION siRNA Universal Negative Control #1 SIC001) at the concentration of 30nM using HiPerFect transfection reagent (Qiagen), according to the manufacturer’s instructions. Briefly, siRNAs and HiPerFect were mixed in serum-free medium and incubated for 10 minutes at room temperature to allow complex formation. The siRNA–HiPerFect complexes were then added dropwise to CAFs plated at 60–70% confluence in complete medium. Cells were harvested 48 to 72 hours post-transfection for downstream analyses. The siRNA sequences are listed in Table S2.

### TERRA silencing with Antisense Oligonucleotides

TERRA was silenced in CAFs using LNA-modified antisense oligonucleotides (ASOs) targeting the telomeric repeat sequence, as previously described (*3*). Briefly, cells were transfected with 100OnM of TERRA-specific ASO or a scrambled control ASO using **HiPerFect** transfection reagent (Qiagen), according to the manufacturer’s protocol. ASOs and HiPerFect were incubated in serum-free medium for 10 minutes at room temperature to allow complex formation, then added dropwise to cells plated at 60–70% confluence in complete medium. Cells were collected 48 hours post-transfection for downstream analyses. The sequences are listed in Table S2.

### Pharmacological Treatments with Small-Molecule Inhibitors

HDFs were treated with the AR antagonist UT-155 (MedChemExpress, Cat. No. HY-12709) at a concentration of 1OµM for 48 hours in medium supplemented with charcoal-stripped fetal bovine serum (FBS). Activated charcoal for serum stripping was obtained from MilliporeSigma. HDFs were also treated with ARCC4 (Tocris, Cat. No. 7254) or its corresponding negative control ARCC4 NC (Tocris, Cat. No. 7255) in charcoal-stripped serum under the same conditions. HDFs and CAFs were treated with the selective androgen receptor modulator Ostarine (MedChemExpress, Cat. No. HY-13273) at 10OµM for 48 hours. Additionally, HDFs and CAFs were treated with the NONO inhibitor (R)-SKBG-1 (MedChemExpress, Cat. No. HY-153918) at 5OµM for either 24 or 48 hours.

### EdU Incorporation Assay

Cell proliferation was assessed using the Click-iT™ EdU Alexa Fluor™ 488 Imaging Kit (Invitrogen), following the manufacturer’s instructions. CAFs were seeded onto 19 mm coverslips in 6-well plates at ∼40% confluence. The following day, cells were incubated with 10OμM EdU for 4 hours, then fixed with 4% paraformaldehyde. Stained samples were imaged using a Zeiss fluorescence microscope. Quantification of EdU-positive cells was performed using ImageJ software, measuring the number of EdU-positive nuclei per field.

### Co-culture Assays

To assess tumor cell expansion in fibroblast–cancer cell co-culture, cells were plated in 8-well chamber slides pre-coated with Matrigel (BD Biosciences). Each well was coated with 100OμL of Matrigel diluted 1:10 in cold medium and incubated at 37O°C for 30 minutes to allow polymerization. HDFs or CAFs were mixed with SCC13 cancer cells at a 1:1 ratio (1,000 fibroblasts and 1,000 cancer cells per well), then seeded onto the coated chambers. Tumor cell expansion was evaluated five days post-seeding by immunofluorescence staining for Pan-keratin (Pan-K, 1:300; Sigma, C2562) to label epithelial cancer cells, Vimentin (Vim, 1:300; Abcam ab137321) to identify fibroblasts, and DAPI (1:1000) to visualize nuclei. Image acquisition was performed using a fluorescence microscope, and quantification of tumor expansion was conducted using ImageJ software.

### Multicellular Spheroid Invasion Assay

Multicellular spheroids were generated from subconfluent cultures using the hanging-drop method (*43*). Briefly, CAFs and/or SCC13 cells were resuspended in complete DMEM supplemented with 10% FBS, 4.8 mg/ml methylcellulose (Sigma-Aldrich, Cat # M6385) and 10 mg/ml bovine dermis collagen I solution (PureCol, Advanced BioMatrix Inc., Cat # 5005,) at a final concentration of 3,000 cells per 30OμL drop. Drops were placed on the inner side of a 10Ocm dish lid and incubated in suspension for 24 hours in an incubator at 37°C and 5% CO_2_ to allow spheroid formation. The pre-formed spheroids were then embedded in 3D collagen matrices. The collagen solution was prepared using mid-density rat-tail collagen (final concentration: 5mg/ml). First, a total volume of 1400 ml, was obtained by adding 112 ml of 10X PBS, 13.89 ml of 1N NaOH, 299.7 ml of 80 mM HEPES in Milli-Q water, and 632.3 ml of non-pepsinized rat-tail collagen type I stock solution (11.07 mg/ml; Corning, Cat #354249) to the mixture. Spheroids composed of CAFs and SCC13 cells (mixed or separate) were washed twice with PBS (5 minutes each), resuspended in 300 ml of medium, and mixed with collagen solution. 150 ml of homogeneous collagen solution was transferred into an 8-chamber slide (at room temperature) with 4-5 spheroids per well. The spheroids were positioned at the interface between collagen and plastic before final polymerization. Invasion has been detected by bright-field microscopy 2 days post-embedding. Image analysis was performed using Fiji/ImageJ. The invasion area was quantified as the 2D area surrounding the spheroid, calculated by subtracting the area of the spheroid core from the total invaded region. Spheroids that fused or were located at the gel edge were excluded from analysis.

### Immunofluorescence Staining and Quantification

For immunofluorescence staining of cultured cells, coverslips were first washed with cold PBS, then fixed with 4% formaldehyde in PBS for 10 minutes at room temperature. Cells were permeabilized with 0.5% Triton X-100 in PBS for 15 minutes and subsequently washed with PBS. Blocking was performed using 2% BSA in PBS for 30 minutes. Cells were then incubated overnight at 4°C with primary antibodies diluted in blocking solution The following day, coverslips were washed three times with PBS and incubated for 1 hour at room temperature with Alexa Fluor-conjugated secondary antibodies (Alexa Fluor 488, 568, or 647; 1:1000 dilution; Thermo Fisher Scientific). After final washes, coverslips were mounted using Dako Fluorescence Mounting Medium (Agilent Technologies). Images were acquired using a Zeiss LSM700 confocal microscope. Quantification of fluorescence signals was performed using Fiji/ImageJ software. For tissue sections, tumors were embedded in OCT compound (Tissue-Tek®), snap-frozen at –80°C, and cryosectioned into 7–8Oμm sections using a cryostat. Sections were air-dried for 30 minutes at room temperature and fixed in 4% paraformaldehyde for 20 minutes. The staining procedure was then carried out as described for cultured cells.

### TERRA RNA FISH

Detection of TERRA RNA foci in HDFs and CAFs was performed using a Stellaris RNA FISH protocol with a Cy3-labeled TelC probe (PNA bio F1002, TelC-Cy3) complementary to the telomeric UUAGGG repeat. Cells were seeded on coverslips and fixed in 4% paraformaldehyde (PFA) in PBS for 10 minutes at room temperature, followed by permeabilization in 70% ethanol at –20°C for at least 1 hour. Hybridization was carried out overnight at 37°C in Stellaris Hybridization Buffer (Biosearch Technologies) supplemented with 125 nM TelC-555 probe in a humidified chamber. For specificity control, parallel samples were treated with RNase A (100Oμg/mL in PBS) for 30 minutes at 37°C prior to hybridization. The following day, coverslips were washed twice for 30 minutes at 37°C in Stellaris Wash Buffer A, counterstained with DAPI, and briefly rinsed in Wash Buffer B. Coverslips were mounted using Prolong Antifade Mounting Medium and imaged using a Nikon CSU-W1 SoRa spinning disk confocal microscope. Quantification of nuclear TERRA foci was performed using Fiji/ImageJ software.

### Immuno-RNA FISH for NONO and TERRA on cells

Combined immunofluorescence and RNA FISH (immuno-RNA FISH) was performed on CAFs to detect the co-localization of the NONO protein and TERRA RNA, using a sequential protocol adapted from Stellaris RNA FISH guidelines. CAFs were seeded on glass coverslips and fixed in 4% paraformaldehyde (PFA) in PBS for 10 minutes at room temperature, followed by permeabilization with 0.5% Triton X-100 in PBS for 10 minutes. After washing, cells were blocked in 2% BSA in PBS for 30 minutes and incubated overnight at 4°C with a rabbit anti-NONO antibody (Proteintech, Cat. No. 10573-1-AP; dilution 1:200) in blocking solution. The next day, cells were washed with PBS and incubated for 1 hour at room temperature with Alexa Fluor-conjugated secondary antibodies (1:1500, Thermo Fisher Scientific). After staining, cells were post-fixed again with 4% PFA for 10 minutes to preserve antibody-antigen complexes during RNA FISH. Following post-fixation, cells were briefly washed in Stellaris Wash Buffer A. Hybridization was performed overnight at 37°C in Stellaris Hybridization Buffer (Biosearch Technologies) containing 125 nM Cy3-labeled TelC probe (PNA bio F1002, TelC-Cy3, complementary to UUAGGG repeats). The following day, cells were washed twice in Stellaris Wash Buffer A for 30 minutes at 37°C, counterstained with DAPI, and briefly rinsed in Wash Buffer B. Coverslips were mounted using Prolong Antifade Mounting Medium and imaged using a Nikon CSU-W1 SoRa spinning disk confocal microscope. Co-localization of TERRA RNA foci and NONO protein was analyzed using Fiji/ImageJ software.

### Immuno-RNA FISH for TERRA and Vimentin in Human Tissue Sections

Immuno-RNA FISH was performed on frozen tissue sections from human SCC and matched adjacent skin and on a frozen tissue array of lung (BioChain, T6235152-5, Lot.C708031) to visualize TERRA RNA and vimentin protein. Sections were air-dried for 30 minutes at room temperature, followed by fixation in 4% paraformaldehyde (PFA) in PBS for 10 minutes. After washing in PBS, sections were permeabilized in 0.5% Triton X-100 in PBS for 10 minutes, then blocked in 5% normal donkey serum (NDS) in PBS for 1 hour at room temperature. Sections were incubated overnight at 4°C with a mouse anti-vimentin antibody (Vim, 1:300; Abcam ab137321) diluted in blocking solution. The next day, sections were washed with PBS and incubated with an Alexa Fluor-conjugated donkey anti-mouse secondary antibody (1:1000, Thermo Fisher Scientific) for 1 hour at room temperature. Following immunostaining, slides were post-fixed in 4% PFA for 10 minutes, then briefly incubated with Stellaris Wash Buffer A. Hybridization was carried out overnight at 37°C in Stellaris Hybridization Buffer (Biosearch Technologies) containing 125 nM TelC-Cy3 probe (PNA bio F1002, TelC-Cy3, complementary to UUAGGG repeats). Post-hybridization, sections were washed twice for 30 minutes at 37°C in Stellaris Wash Buffer A, counterstained with DAPI, and briefly rinsed in Wash Buffer B. Slides were mounted using Prolong Antifade Mounting Medium and imaged using a Nikon CSU-W1 SoRa spinning disk confocal microscope. The percentage of vimentin-positive cells in which at least one TERRA focus has been detected was analyzed using Fiji/ImageJ software.

### Proximity Ligation Assay (PLA)

Proximity ligation assays (PLAs) were performed using the Duolink® In Situ Red Starter Kit (Sigma-Aldrich, Cat. No. DUO92101) according to the manufacturer’s instructions. Cells were seeded on glass coverslips in a 24-well plate and fixed in cold 4% paraformaldehyde (PFA) for 15 minutes at room temperature, followed by three washes with PBS. After fixation, cells were permeabilized with 0.1% Triton X-100 in PBS for 15 minutes at room temperature and then incubated with Duolink Blocking Solution for 1 hour at 37°C in a humidified chamber. Following blocking, cells were incubated overnight at 4°C with primary antibodies diluted in Duolink Antibody Diluent. The next day, cells were washed and incubated with the appropriate Duolink PLA probes (PLUS and MINUS) for 1 hour at 37°C in a humidified chamber. After washing three times with Buffer A (provided in the kit), cells were incubated with the ligation mix for 1 hour at 37°C, followed by another three washes in Buffer A. Amplification was then performed by incubating the cells with the amplification mix for 140 minutes at 37°C in a dark, humidified chamber. After amplification, cells were washed twice in Buffer B for 10 minutes each, followed by a brief wash in 0.01× Buffer B for 1 minute. Coverslips were mounted with Duolink In Situ Mounting Medium containing DAPI. Images were acquired using a Zeiss microscope, and PLA signals were quantified using ImageJ software by performing fluorescent particle analysis to count the number of PLA puncta per nucleus per field.

Antibodies used were anti-mouse TRF2 monoclonal antibody (Cat. 13579, Abcam, 1:50 dilution), anti-rabbit Phospho-Histone H2A.X (Ser139) (20E3) (Cell Signalling #9718 1:100), anti-rabbit NONO (Proteintech Cat No. 11058-1-AP, 1:200) and anti-mouse AR (Santa Cruz sc-7305, 1:100).

### Immunoblotting

Cells were lysed directly in NuPAGE™ LDS Sample Buffer (4X; Thermo Fisher Scientific, NP0007) supplemented with 1% β-mercaptoethanol and protease/phosphatase inhibitors (Thermo Fisher Scientific). Lysates were collected and briefly sonicated, then boiled at 95°C for 5 minutes to ensure complete protein denaturation. An equal volume of protein was loaded and resolved on 4–12% Bis-Tris NuPAGE™ gels (Thermo Fisher Scientific) using MOPS SDS Running Buffer and subsequently transferred to PVDF membranes using the iBlot™ 2 Dry Blotting System (Thermo Fisher Scientific). Membranes were blocked in 5% non-fat dry milk in TBS-T (Tris-buffered saline with 0.1% Tween-20) for 1 hour at room temperature and incubated overnight at 4°C with primary antibodies diluted in blocking buffer. After washing, membranes were incubated with HRP-conjugated secondary antibodies for 1 hour at room temperature. Protein bands were visualized using enhanced chemiluminescence (ECL; Thermo Fisher Scientific) and imaged with Azure Biosystem Chemidoc Imaging System or detected on Fuji Medical X-ray films (Fujifilm).

### Gene Expression Analysis by Quantitative Real-Time PCR

Total RNA was extracted using the Direct-zol™ RNA Miniprep Kit (Zymo Research, Cat# R2050) according to the manufacturer’s instructions, including on-column DNase I treatment to eliminate genomic DNA contamination. RNA concentration and purity were assessed using a NanoDrop spectrophotometer (Thermo Fisher Scientific). For cDNA synthesis, 500Ong of total RNA was reverse transcribed using the iScript™ Reverse Transcription Supermix (Bio-Rad, Cat# 1708840) in a 20OµL reaction, following the manufacturer’s protocol. Quantitative real-time PCR (qPCR) was performed on an Applied Biosystems QuantStudio™ 5 Real-Time PCR System using Applied Biosystems SYBR™ Green PCR Master Mix (Cat# 4309155). Gene expression was analyzed using the comparative Ct (ΔΔCt) method. All reactions were run in technical triplicates, and expression levels were normalized to the endogenous control gene RPLP0.

### RNA Extraction, DNase Treatment, Reverse Transcription, and RT-qPCR for TERRA

Total RNA was extracted from cultured cells using the Direct-zol™ RNA Miniprep Kit (Zymo Research, Cat# R2050), following the manufacturer’s instructions. Cells were lysed in TRIzol™ reagent and processed through Zymo-Spin™ columns, including successive on-column DNase I treatment to ensure thorough removal of genomic DNA. RNA was eluted in RNase-free water and quantified using a NanoDrop spectrophotometer (Thermo Fisher Scientific). Only samples with OD260/280 ≥ 1.8 were used for downstream applications.

Since only approximately 7% of TERRA transcripts are polyadenylated, the majority do not undergo full polyA maturation (*44*). To capture the broad population of TERRA transcripts, reverse transcription was performed using a TERRA-specific primer composed of telomeric repeats (Supplementary Table 2). To enable normalization, a GAPDH gene-specific reverse primer was included in the same reverse transcription reaction. (Supplementary Table 2). Reverse transcription was carried out using 1Oµg of total DNase-treated RNA (as previously described), combined with the iScript™ Reverse Transcription Supermix (Bio-Rad, Cat# 1708840) in a 20 µL final volume and following the manufacturer’s protocol. A no-reverse transcriptase (–RT) control was included to confirm the absence of genomic DNA contamination. The resulting cDNA was used for quantitative PCR (RT-qPCR) using specific primer pairs targeting TERRA transcripts from subtelomeric regions (Supplementary Table 2) and GAPDH as the internal normalization control. All reactions were performed in technical triplicate and analyzed using the ΔΔCt method.

### RNA Immunoprecipitation (RIP) for TERRA Enrichment in CAFs and HDFs

RNA immunoprecipitation (RIP) was performed in CAFs or HDFs to assess the interaction between the RNA-binding protein NONO and TERRA transcripts originating from chromosome 7p. Approximately 4 million cells per condition were harvested, washed with cold PBS, and lysed in RIP lysis buffer (10 mM HEPES pH 7.4, 5 mM MgCl2,100 mM KCl, 2 mM EDTA pH 8, 0.5% NP-40, 0.5 mM DTT), supplemented with RNase and protease inhibitors. Lysates were cleared by centrifugation at 12,000 × g for 10 minutes at 4°C, precleared with anti-rabbit IgG magnetic beads (Invitrogen, Cat# 11204D) for 1 hour at 4°C, and incubated for 1 hour at RT with 5 µg of anti-NONO antibody (Proteintech, Cat# 22646-1-AP) or 5 µg of normal rabbit IgG (control). Immune complexes were captured by incubation with anti-rabbit IgG magnetic beads for 3 hours at 4°C under rotation. Beads were washed extensively with high-salt wash buffer (100 mM NaCl, 10 mM Tris-HCl pH 7.4, 0.5% SDS, 1 mM EDTA pH 8) containing RNase and protease inhibitors. RNA was extracted from immunoprecipitates using TRIzol reagent (Invitrogen), precipitated with ethanol, and reverse transcribed as previously described. Quantitative PCR (RT-qPCR) was performed using primers specific for TERRA transcripts from chromosome 7p. Amplification of RPLP0 mRNA was included as a negative control for NONO binding. A 10% input sample was processed in parallel and used for normalization.

Enrichment was calculated using the percentage of input method: %Input=100×2(CtInput−CtIP).

### Chromatin Immunoprecipitation (ChIP) and qPCR

Primary human dermal fibroblasts (HDFs) with or without AR silencing were cross-linked with formaldehyde at a final concentration of 1% for 10 minutes at room temperature to preserve protein–DNA interactions. The reaction was quenched by adding glycine to a final concentration of 125 mM. Cells were washed with ice-cold PBS and collected by centrifugation at 400 × g. Cell lysis and chromatin preparation were performed using the iDeal ChIP-seq Kit for Transcription Factors (Diagenode) according to the manufacturer’s instructions. Chromatin was fragmented to a size range of 100–300 bp using a Diagenode Bioruptor sonicator. Samples were precleared using magnetic beads provided in the kit and incubated overnight at 4°C with 5 µg of anti-Androgen Receptor (AR) antibody (Cell Signaling Technology, clone D6F11) or with an equivalent amount of non-immune rabbit IgG (Diagenode) as control. Immunocomplexes were pulled down using DiaMag protein A-coated magnetic beads (Diagenode). Elution, reverse crosslinking, and DNA purification were carried out following the kit protocol. Immunoprecipitated DNA was quantified using the Qubit Fluorometric Quantification Kit (Thermo Fisher Scientific). Enrichment at target regions was assessed by qPCR using SYBR Green-based detection (Applied Biosystems) and expressed as percent input. Primer sequences targeting AR sites on TERRA subtelomeric promoter regions are listed in Supplementary Table S2.

### Identification of TERRA-Interacting Proteins by iDRIP

iDRIP (identification of direct RNA-interacting proteins) was performed in cancer-associated fibroblasts (CAFs) following the protocol by Chu et al, with minor modifications (*3*). Approximately 10 million cells per condition were UV-crosslinked at 254 nm (400 mJ/cm²) to preserve endogenous RNA-protein interactions. Cells were lysed in iDRIP lysis buffer (20 mM Tris-HCl pH 7.5, 150 mM NaCl, 1 mM EDTA, 1% NP-40, 0.1% SDS, 0.5% sodium deoxycholate, 1 mM DTT), supplemented with protease and RNase inhibitors. Lysates were treated with DNase and low concentrations of RNase I (Thermo Fisher Scientific) to fragment RNA while preserving RNA-protein crosslinks. Pulldown was carried out using a biotinylated antisense DNA probe complementary to the UUAGGG repeat region of TERRA (TERRA-AS), or a biotinylated sense probe (TERRA-SENSE) was used in parallel as a negative control, hybridized at 37°C for 2 hours. Streptavidin-conjugated magnetic beads were used to isolate probe-bound complexes and sent to the Harvard Taplin Biological Mass Spectrometry facility for LC-MS/MS.

### Bead Digestion Analysis by LC-MS/MS

Beads were washed at least five times with 100ul 50 mM ammonium bicarbonate, then 5ul (200ng/ul) of modified sequencing-grade trypsin (Promega, Madison, WI) was spiked in and the samples were placed in a 37°C room overnight. The samples were then centrifuged or placed on a magnetic plate if magnetic beads were used, and the liquid was removed. The extracts were then dried in a speed-vac (∼1 hr). Samples were then re-suspended in 50ul of HPLC solvent A (2.5% acetonitrile, 0.1% formic acid) and desalted by STAGE tip (*45*). On the day of analysis, the samples were reconstituted in 10 µl of HPLC solvent A. A nano-scale reverse-phase HPLC capillary column was created by packing 2.6 µm C18 spherical silica beads into a fused silica capillary (100 µm inner diameter x ∼30 cm length) with a flame-drawn tip (*46*). After equilibrating the column, each sample was loaded via a Famos auto sampler (LC Packings, San Francisco, CA) onto the column. A gradient was formed and peptides were eluted with increasing concentrations of solvent B (97.5% acetonitrile, 0.1% formic acid). As peptides eluted, they were subjected to electrospray ionization and then entered into a Velos Orbitrap Elite ion-trap mass spectrometer (Thermo Fisher Scientific, Waltham, MA). Peptides were detected, isolated, and fragmented to produce a tandem mass spectrum of specific fragment ions for each peptide. Peptide sequences (and hence protein identity) were determined by matching protein databases with the acquired fragmentation pattern by the software program, Sequest (Thermo Fisher Scientific, Waltham, MA) (*47*). All databases include a reversed version of all the sequences, and the data was filtered to a one and two-percent peptide false discovery rate.

### Transcriptomic Analysis

Total RNA was extracted and purified using the Direct-zol RNA Miniprep Kit (Zymo Research). RNA quality was assessed to ensure OD260/OD280 ≥ 1.8 and RNA Integrity Number (RIN) ≥ 8. The GeneChip® WT PLUS Reagent Kit (Thermo Fisher Scientific) was used for sample preparation, and hybridization was performed on the human Clariom™ D Arrays (Thermo Fisher Scientific). Microarray experiments and data acquisition were carried out at the Institute of Genetics and Genomics of Geneva (iGE3). Data processing and analysis were performed using the Transcriptome Analysis Console (TAC) software (Thermo Fisher Scientific). Global transcriptomic changes were further analyzed using Gene Set Enrichment Analysis (GSEA) with curated gene signatures from the MSigDB v7.2 database (https://software.broadinstitute.org/cancer/software/gsea).

### Intradermal Back Injection in Mice

Intradermal tumorigenicity assays were performed in 6- to 10-week-old male and female NOD/SCID mice (Jackson Laboratory). A total of 2.5 × 10O SCC13 cells were admixed with an equal number of human dermal fibroblasts (HDFs) with or without TERRA overexpression, or with TERRA-silenced cancer-associated fibroblasts (CAFs). After centrifugation, cell pellets were resuspended in 100 µL of Matrigel (BD Biosciences) and injected intradermally into the left and right flanks of the mouse back. Mice were sacrificed 10 days post-injection for tissue analysis. Tumor volume was calculated using the formula: V *(Length × Width²) / 2*, where V is volume, W is width, and L is length. Tumors did not exceed the maximum dimension permitted by the approved protocol (20 mm in any direction). For intratumoral delivery of (R)-SKBG-1, a total of 2.5 × 10O SCC13 cells were admixed with an equal number of CAFs. Upon detection of palpable lesions (∼5 mm in diameter) 7 days post-injection, (R)-SKBG-1 (10 µM diluted in PBS) or vehicle (PBS) was administered intratumorally into paired lesions once daily for 14 days. All animal procedures were conducted in accordance with Massachusetts General Hospital Institutional Animal Care and Use Committee (IACUC)-approved protocol (2004N000170).

### Ex vivo Biopsy Culture and Treatment

Freshly collected surplus biopsies from cutaneous squamous cell carcinoma (cSCC), actinic keratosis, and lentigo maligna melanoma were bisected and immediately used for ex vivo culture as described before(*48*). Briefly, the biopsies were placed in cell culture inserts (Millipore, Cat. No. Z352985). Inserts were positioned in 6-well plates containing 1.5OmL of co-culture medium in the lower compartment only. Biopsies were treated with either DMSO (vehicle control) or (R)-SKBG1 (10OµM) and maintained for 2 days, with daily medium replacement. Following the treatment period, hematoxylin and eosin (H&E) staining was performed to assess tissue integrity. The co-culture medium consisted of 3 parts DMEM (Gibco, Cat. No. 41966052) and 1 part Ham’s F-12 Nutrient Mix (Gibco, Cat. No. 11765054), supplemented with 10% heat-inactivated fetal bovine serum (Gibco, Cat. No. 1050), 0.1Omg/mL Normocin™ (Invivogen, Cat. No. ant-nr-1), 21.8Oμg/mL adenine (Sigma-Aldrich, Cat. No. A2786), 5.45Oμg/mL apotransferrin (Sigma-Aldrich, Cat. No. T1147), 2.18OnM triiodothyronine (Sigma-Aldrich, Cat. No. T6397), 0.44Oμg/mL hydrocortisone (Sigma-Aldrich, Cat. No. H0888), 0.11OnM cholera toxin (Sigma-Aldrich, Cat. No. C8052), 5.50Oμg/mL insulin (Sigma-Aldrich, Cat. No. I6634), and 0.01Oμg/mL epidermal growth factor (Sigma-Aldrich, Cat. No. E4127).

### Statistical significance

All statistical analyses were performed using GraphPad Prism version 10.5.0 (GraphPad Software, Inc.). As indicated in the figure legends, data are presented as mean ± standard deviation (SD). Detailed descriptions of the statistical methods applied to each experiment are provided in the corresponding legends. Unless otherwise noted, two-tailed Student’s t-tests were used to compare two groups. For comparisons involving more than two groups, one-way ANOVA was performed, followed by Sidak’s, Tukey’s, or Dunnett’s multiple comparison tests, as appropriate.

### Language Editing

Large language models, including ChatGPT (OpenAI) were employed to assist in improving the clarity and fluency of English in the manuscript text. These tools were not used for data analysis, interpretation, or generation of scientific content.

## Data Availability

Raw and processed datasets used for this article are available under the repository accession numbers GSE300351 (Transcriptomic analysis of CAFs with silencing of NONO); GSE300353 (Transcriptomic analysis of CAFs with silencing of TERRA lncRNA

## Acknowledgements

We thank the Molecular Imaging Core at MGH, led by Dr. Roy Soberman.

## Fundings

This study was supported by grants from the NIH (R01AR039190, R01AR078374, R01CA269356; the contents are solely the responsibility of the authors and do not necessarily represent the official views of the NIH). E.D.C. was supported by a fellowship from the Italian Association for Cancer Research (AIRC) (2020A009262) and the Dermatology Foundation Research Grant (2024A014581). J.I. received support from the European Union’s Horizon 2020 Research and Innovation Programme under the Marie Skłodowska-Curie Grant Agreement No. 859860. G.P.D. is a member of the SKINTEGRITY.CH collaborative research program.

## Authors contribution

EDC, JI, SS, and AK performed experiments and analyzed the results with GPD. AM provided the reagents for the RIP and spheroids assay and contributed to the proofreading of the manuscript. SG contributed to the proofreading of the manuscript. EDC and JI performed the bioinformatic analysis of transcriptomics with GPD. VN provided the human specimens. ED and GPD designed the study and wrote the manuscript.

## Competing interests

The authors declare the following financial interests, which may be considered potential competing interests: GPD is a co-founder (with equity) of the EpiKare International company. AM is a co-founder (with equity) of New Frontier Bio, a consumer health company developing skincare and anti-aging products, and has equity in DermBiont, a private company advancing targeted topical therapeutics for dermatologic indications. AM and SS receive research support from Shiseido Inc. The other authors declare no competing interests.

