## Supplemental Figures and Figure legends for "A TERRA–NONO Axis Drives Fibroblast Reprogramming in Cancer"

#### Supplementary Figures and figure legends

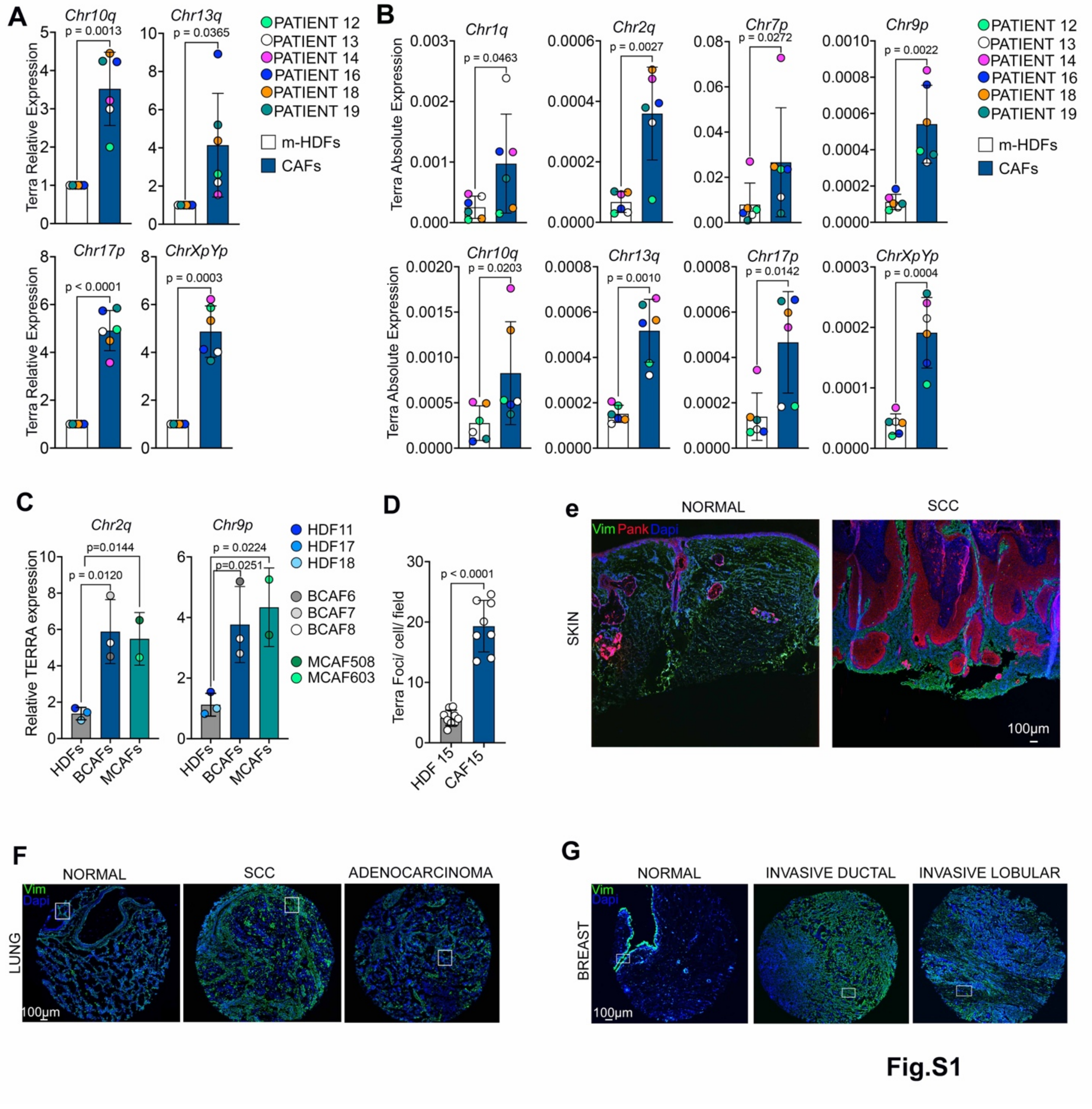


**fig.S1. Terra is highly expressed in CAFs vs matched fibroblasts.**

**(A)** Chromosome-specific TERRA expression in multiple CAF strains isolated from skin SCC lesions (CAFs) compared to HDFs from adjacent unaffected skin (HDFs) from the same patients as in Fig.1a, as determined by RT-qPCR using specific primers. Expression is shown as fold change in CAFs versus matched HDFs, normalized to GAPDH. n = 6 (strains). Mean ± SD; two-tailed paired t-test.

**(B)** Absolute TERRA expression in multiple CAF compared to dermal fibroblasts from adjacent unaffected skin (HDFs) of the same patients, assessed by RT-qPCR as in Figure 1a. mean ± SD. Two-tailed paired *t*-test.

**(C)** RT-qPCR analysis of chromosome-specific TERRA expression in CAFs isolated from BCC lesions (BCAFs) (BCAF6, BCAF7, BCAF8) and melanomas (MCAF508, MCAF603) compared to three age-matched reference HDFs (HDF11, HDF17, HDF18) normalized to GAPDH. n = 3 strains (BCAFs) n=2 strains (MCAFs). Mean ± SD. One-way ANOVA with Dunnett's multiple comparisons test.

**(D)** Representative images of TERRA FISH (red) and quantification of TERRA nuclear foci per cell, averaged per field, in an additional CAF strain compared to matched HDFs from the same patients, as in Fig. 1C. n = 8 to 11 fields per sample. Mean ± SD. Unpaired t-test with Welch's correction.

**(E)** Representative immunofluorescence images of normal skin and squamous cell carcinoma (SCC) sections stained for vimentin (Vim; green), pan-keratin (Pank; red), and DAPI (blue) to distinguish stromal and epithelial compartments. These samples correspond to SCC lesions subsequently analyzed by combined TERRA RNA FISH and immunofluorescence in parallel sections.

**(F)** Low magnification images of normal lung tissue, lung squamous cell carcinoma (SCC), and lung adenocarcinoma from a tissue array stained with anti-vimentin antibody to identify stromal cells and TERRA RNA FISH. The high magnification area shown in Fig.1e is indicated by a square.

**(G)** Low magnification images of normal breast tissue, invasive ductal carcinoma, and invasive lobular carcinoma from a tissue array stained with anti-vimentin antibody to identify stromal cells and TERRA RNA FISH. The high magnification area shown in Fig.1f is indicated by a square.

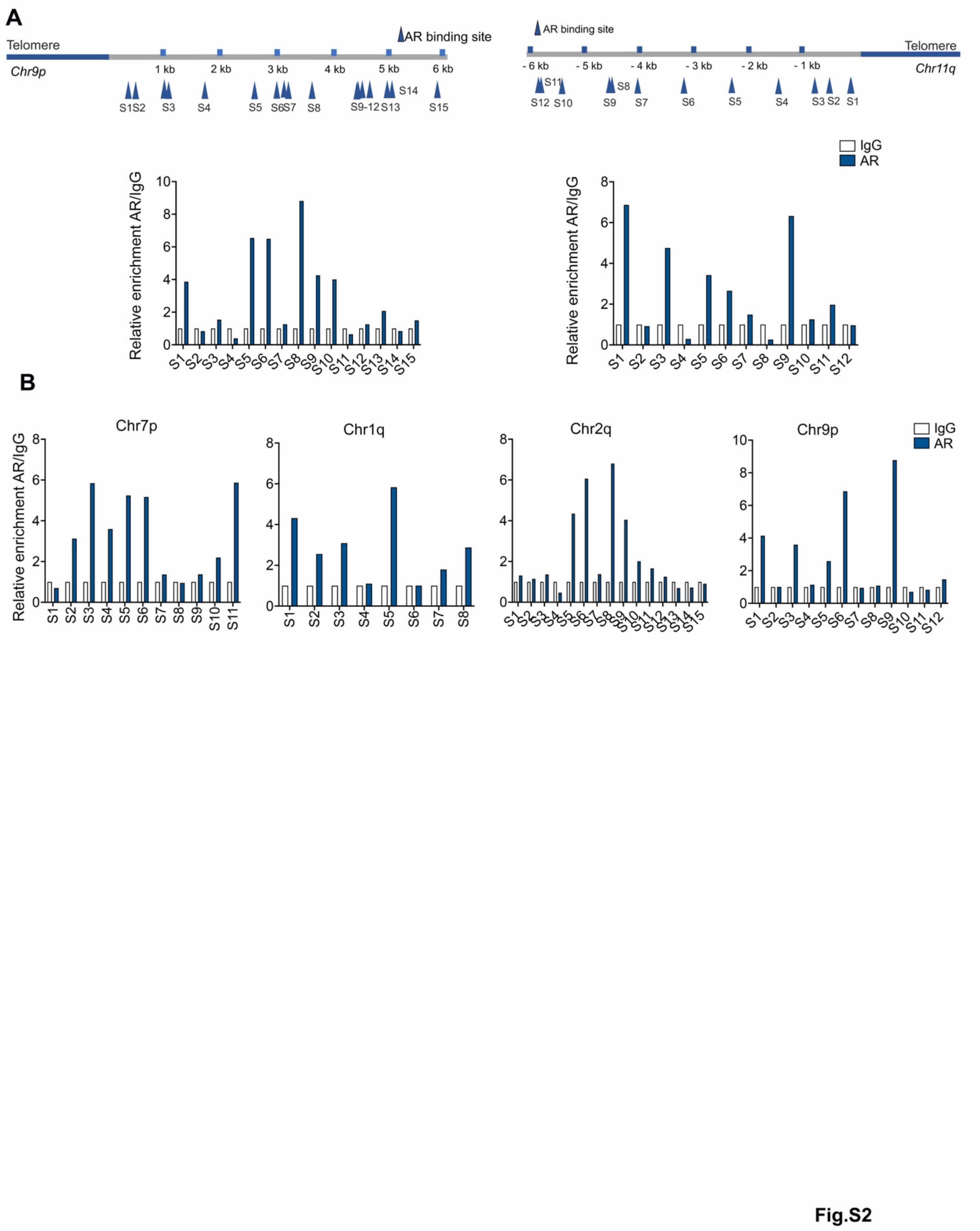


**Fig.S2. AR binds to subtelomeric regions in HDFs**

**(A)** Top: Map of predicted AR binding sites (blue arrowheads) within the sub-telomeric 6 kb regions of the indicated chromosomes. Bottom: ChIP-qPCR analysis of AR binding at these sites in HDF1 using AR antibodies or nonimmune IgG controls. Data are expressed as fold enrichment over IgG controls.

**(B)** ChIP-qPCR analysis of AR binding at these sites in HDF2 using AR antibodies or nonimmune IgG controls. Data are expressed as fold enrichment over IgG controls.

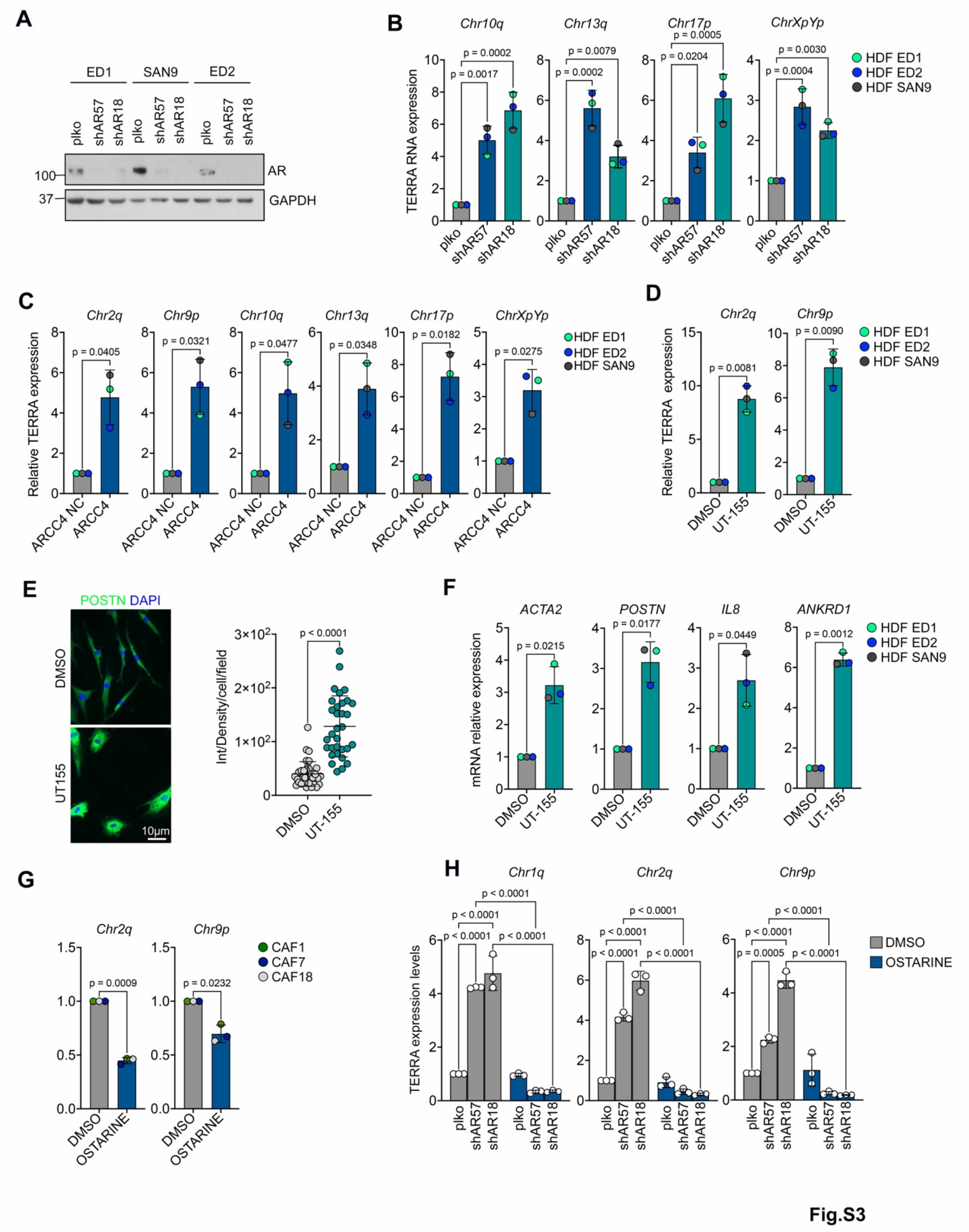


**Fig. S3. TERRA Transcription Is Repressed by AR in HDFs**

**(A)** Immunoblot analysis with anti-AR and GAPDH antibodies of three HDFs strains following AR silencing by infection with two different shRNAs silencing lentiviruses (shAR#57, shAR#18) versus control (plko).

**(B)**Chromosome-specific TERRA expression in multiple HDF strains following AR silencing by infection with two different shRNAs silencing lentiviruses (shAR#57, shAR#18) versus control (plko), measured by RT-qPCR with specific primers. Expression is shown as fold change relative to control, normalized to GAPDH. n = 3 HDF strains. Data represent mean ± SD. One-way ANOVA with Dunnett's multiple comparisons test.

**(C)** Chromosome-specific TERRA expression (in three HDF strains treated with ARCC4 (1 µM, 48 h) versus ARCC4-negative control (ARCC4 NC) as in the previous panel. Expression is shown as fold change relative to ARCC4 NC, normalized to GAPDH. n = 3 strains. Mean ± SD. Paired two-tailed t-test.

**(D)** Chromosome-specific TERRA expression in three HDF strains treated with UT-155 (1 µM, 48 h) versus DMSO control, expressed as fold change relative to DMSO. n = 3 strains. Mean ± SD. Paired two-tailed t-test.

**(E)** Representative images and quantification of immunofluorescence with antibody against periostin (POSTN) (green) in HDFs treated with UT-155 versus DMSO control as described before. Quantification represents the integrated density of fluorescence per cell per field. Data represent pooled fields from 3 independent experiments (n>30 fields total). Mean ± SD. Unpaired t-test with Welch's correction.

**(F)** Expression of the indicated CAF effector genes in HDFs treated with UT-155 versus DMSO, as in the previous panel, assessed by RT-qPCR. Data are expressed as fold change relative to DMSO. n=3 CAF strains. Mean ± SD. Two-tailed paired t-test.

**(G)** Chromosome-specific TERRA expression in multiple CAF strains treated with Ostarine (10 µM, 48 h) versus DMSO. Results are expressed as fold changes relative to DMSO. n = 3 strains. Mean ± SD. Two-tailed paired t-test.

**(H)** RT-qPCR analysis of TERRA expression (Chr1q, Chr2q, Chr9p) in HDFs after AR silencing (shAR#57, shAR#18) versus control (plko) treated with ostarine (10 µM, 48 h) or DMSO as in the previous panel. Data are expressed as fold change relative to plko with DMSO. Mean ± SD. n = 3 biological replicates. One-way ANOVA with Šídák's multiple comparisons test.

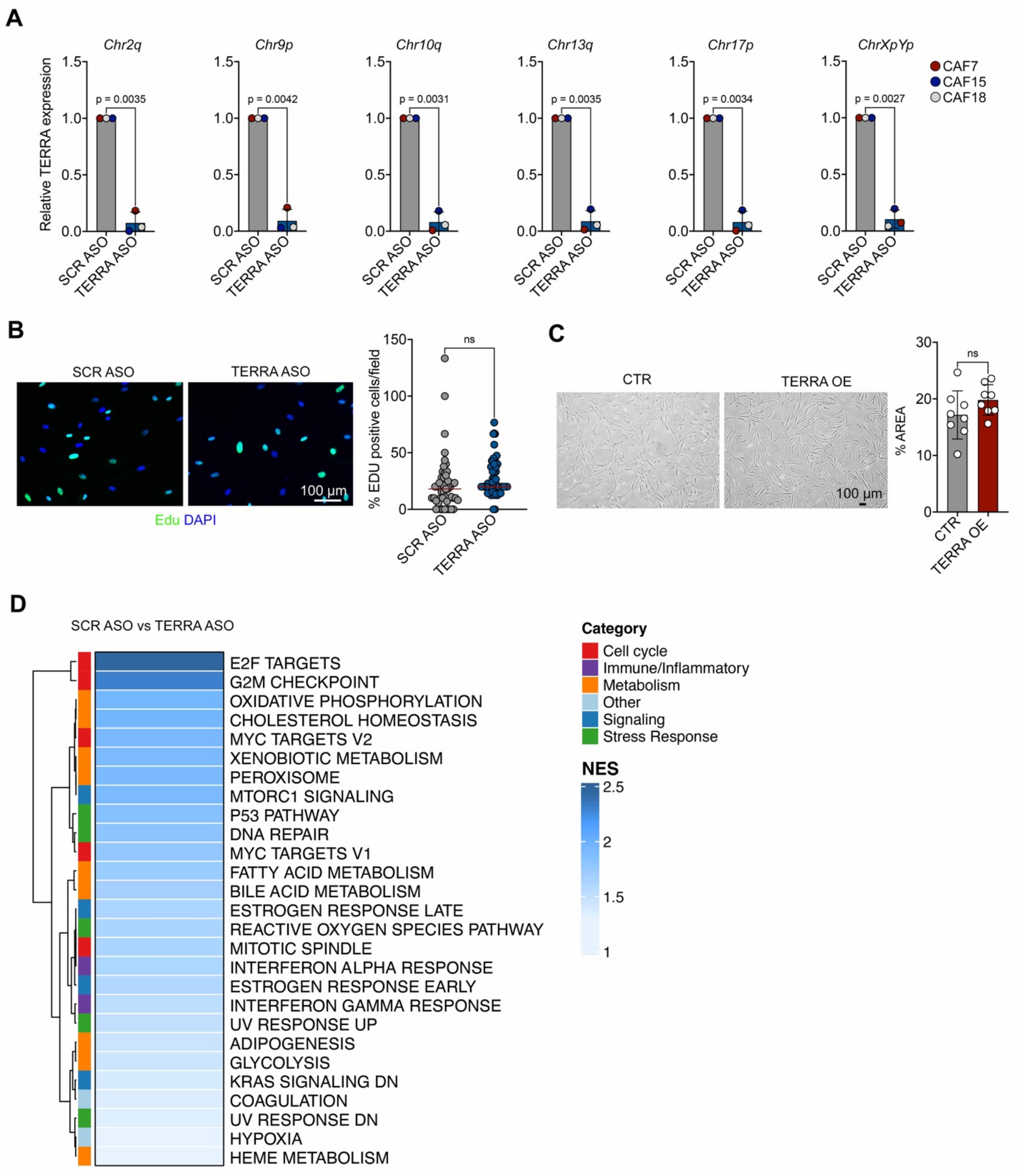
A
Chr2q
Chr9p
Chr10q
Chr13q
Chr17p
ChrXpYp
Relative TERRA expression
p = 0.0035
p = 0.0042
p = 0.0031
p = 0.0035
p = 0.0034
p = 0.0027
SCR ASO
TERRA ASO
CAF7
CAF15
CAF18
B
SCR ASO
TERRA ASO
100 µm
Edu DAPI
% EDU positive cells/field
ns
C
CTR
TERRA OE
100 µm
% AREA
ns
D
SCR ASO vs TERRA ASO
E2F TARGETS
G2M CHECKPOINT
OXIDATIVE PHOSPHORYLATION
CHOLESTEROL HOMEOSTASIS
MYC TARGETS V2
XENOBIOTIC METABOLISM
PEROXISOME
MTORC1 SIGNALING
P53 PATHWAY
DNA REPAIR
MYC TARGETS V1
FATTY ACID METABOLISM
BILE ACID METABOLISM
ESTROGEN RESPONSE LATE
REACTIVE OXYGEN SPECIES PATHWAY
MITOTIC SPINDLE
INTERFERON ALPHA RESPONSE
ESTROGEN RESPONSE EARLY
INTERFERON GAMMA RESPONSE
UV RESPONSE UP
ADIPOGENESIS
GLYCOLYSIS
KRAS SIGNALING DN
COAGULATION
UV RESPONSE DN
HYPOXIA
HEME METABOLISM
Category
Cell cycle
Immune/Inflammatory
Metabolism
Other
Signaling
Stress Response
NES

**Fig.S4**

**Fig. S4. Effect of TERRA Silencing and Overexpression on CAF Proliferation and Gene Expression Programs**

**(A)** Chromosome-specific TERRA expression in CAFs 48 hours after transfection with TERRA ASO or SCR ASO (100 nM). Data are expressed as relative fold change after GAPDH normalization. n=3 strains. Mean ± SD. Two-tailed paired t-test.

**(B)** Proliferation of CAF transfected with TERRA ASO or SCR ASO assessed by EdU incorporation assay (green). Quantification shows the number of Edu-positive cells per field. Data represent pooled fields from 3 independent experiments (n>40 fields total). Mean ± SD. Unpaired t-test with Welch's correction. p=0.1194.

**(C)** Bright-field images of the cell density of HDFs infected with TERRA OE lentiviral vector or CTR vector to assess the cellular proliferation. Quantification shows the percentage of area covered by the cells. Data represent pooled fields from 3 independent experiments (n=8 fields total). Mean ± SD. Unpaired t-test with Welch's correction. p=0.1645.

**(D)** Gene Set Enrichment Analysis (GSEA) of the 50 Hallmark gene sets from MSigDB (https://www.gsea-msigdb.org/gsea/msigdb/), comparing CAFs treated with SCR ASO versus TERRA ASO. Shown are the gene sets significantly enriched in SCR ASO, ranked by Normalized Enrichment Score (NES). Functional categories are color-coded: cell cycle (red), immune/inflammatory (purple), metabolism (orange), signaling (blue), stress response (green), and other (gray). NES values are represented by the gradient scale on the right.

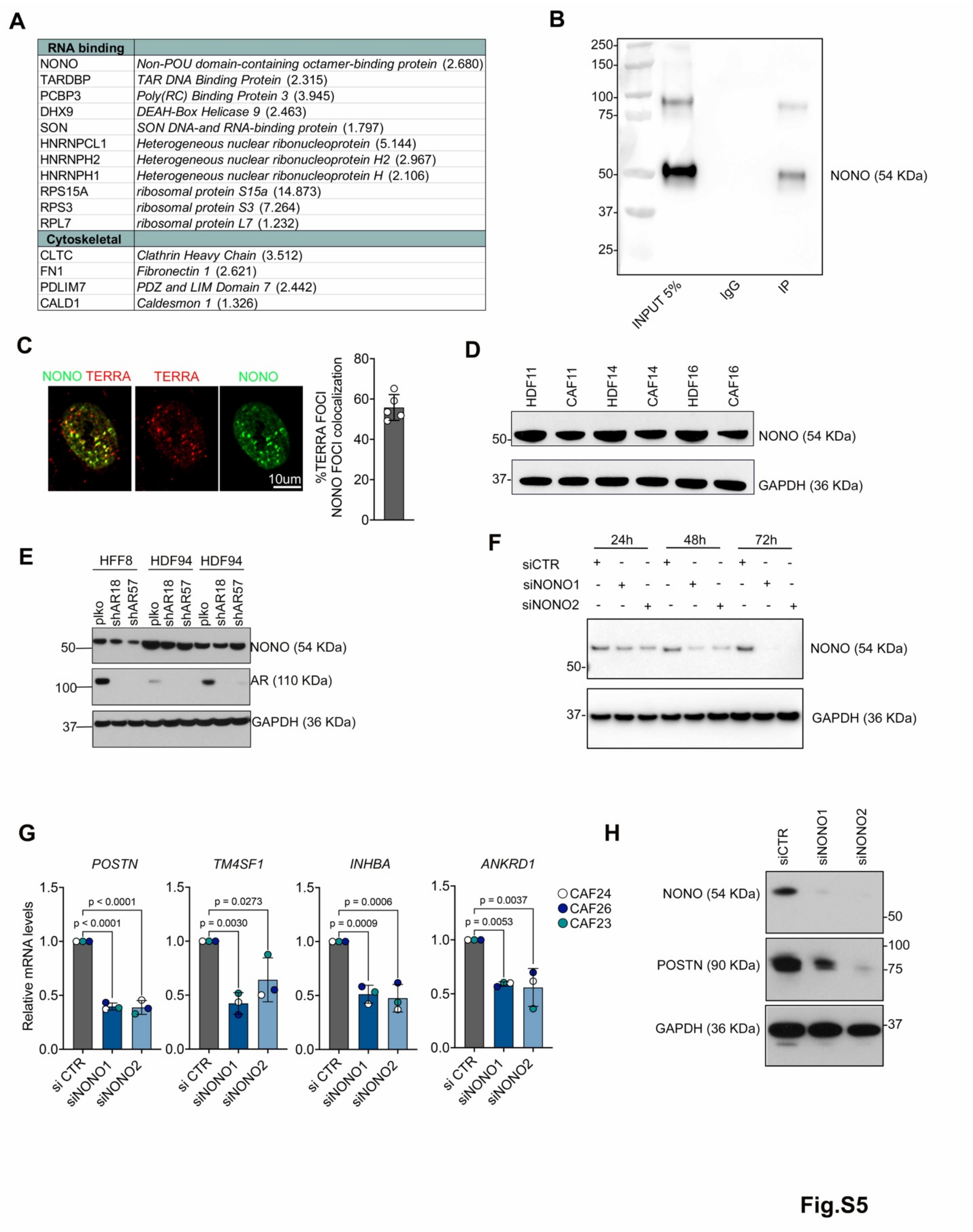

| RNA binding | |
|---|---|
| NONO | *Non-POU domain-containing octamer-binding protein* (2.680) |
| TARDBP | *TAR DNA Binding Protein* (2.315) |
| PCBP3 | *Poly(RC) Binding Protein 3* (3.945) |
| DHX9 | *DEAH-Box Helicase 9* (2.463) |
| SON | *SON DNA-and RNA-binding protein* (1.797) |
| HNRNPCL1 | *Heterogeneous nuclear ribonucleoprotein* (5.144) |
| HNRNPH2 | *Heterogeneous nuclear ribonucleoprotein H2* (2.967) |
| HNRNPH1 | *Heterogeneous nuclear ribonucleoprotein H* (2.106) |
| RPS15A | *ribosomal protein S15a* (14.873) |
| RPS3 | *ribosomal protein S3* (7.264) |
| RPL7 | *ribosomal protein L7* (1.232) |
| **Cytoskeletal** | |
| CLTC | *Clathrin Heavy Chain* (3.512) |
| FN1 | *Fibronectin 1* (2.621) |
| PDLIM7 | *PDZ and LIM Domain 7* (2.442) |
| CALD1 | *Caldesmon 1* (1.326) |



### Fig. S5 Expression and Regulation of NONO in CAFs and HDFs

**(A)** List of TERRA-interacting proteins commonly identified in two independent biological replicates of iDRIP-MS performed on CAFs (strain #19) following pulldown with a biotinylated TERRA antisense (TERRA-AS) oligonucleotide, compared to a negative control (TERRA-sense). Proteins were selected based on their detection in both replicates and filtered for $Log_2$ fold enrichment of TERRA-AS over TERRA-sense ≥ 1, calculated from the normalized intensity values of recovered proteins. Fold enrichment values are reported in brackets.

**(B)** Immunoblot analysis of CAF lysates following immunoprecipitation with anti-NONO antibody or non-immune IgG control. The blot was probed with an anti-NONO antibody. A specific band corresponding to NONO (54 kDa) is detected in the input and specifically enriched in the NONO IP lane, with no signal in the IgG control, confirming the specificity of the immunoprecipitation.

**(C)** TERRA-NONO association in CAFs as detected by immuno-RNA FISH. Shown are representative images of TERRA (red) and NONO (green) foci, detected by RNA FISH with a TERRA-specific probe (red) and IF with anti-NONO antibodies (green), together with a quantification of the percentage of TERRA/NONO colocalizing foci (yellow) per cell. Images were acquired by SoRa super-resolution microscopy with a 60× objective. n = 5 fields (with a minimum of 10 cells per field).

**(D)** Immunoblot analysis with anti-NONO and GAPDH antibodies of CAFs compared to matched HDFs. n=3 strains.

**(E)** Immunoblot analysis with anti-NONO, anti-AR, and GAPDH antibodies of HDFs after AR silencing (shAR#57, shAR#18) versus control (plko). n=3 strains

**(F)** Immunoblot analysis with anti-NONO and GAPDH antibodies of CAFs at different time points (hours) after transfection with two NONO targeting siRNAs (siNONO#1, siNONO#2) versus control (siCTR).

**(G)** Expression of the indicated CAF effector genes (POSTN, TM4SF1, INHBA, ANKRD1) assessed by RT-qPCR in three CAF strains with or without NONO silencing, as described above. Data are expressed as fold change relative to siCTR and normalized to RPLP0. n = 3 strains. Mean ± SD. One-way ANOVA with Dunnett's multiple comparison correction.

**(H)** Immunoblot analysis of CAFs (CAF19) 72 hours after transfection with control or two different NONO-targeting siRNAs. Blots were probed with antibodies against NONO, periostin (POSTN), and GAPDH (loading control).

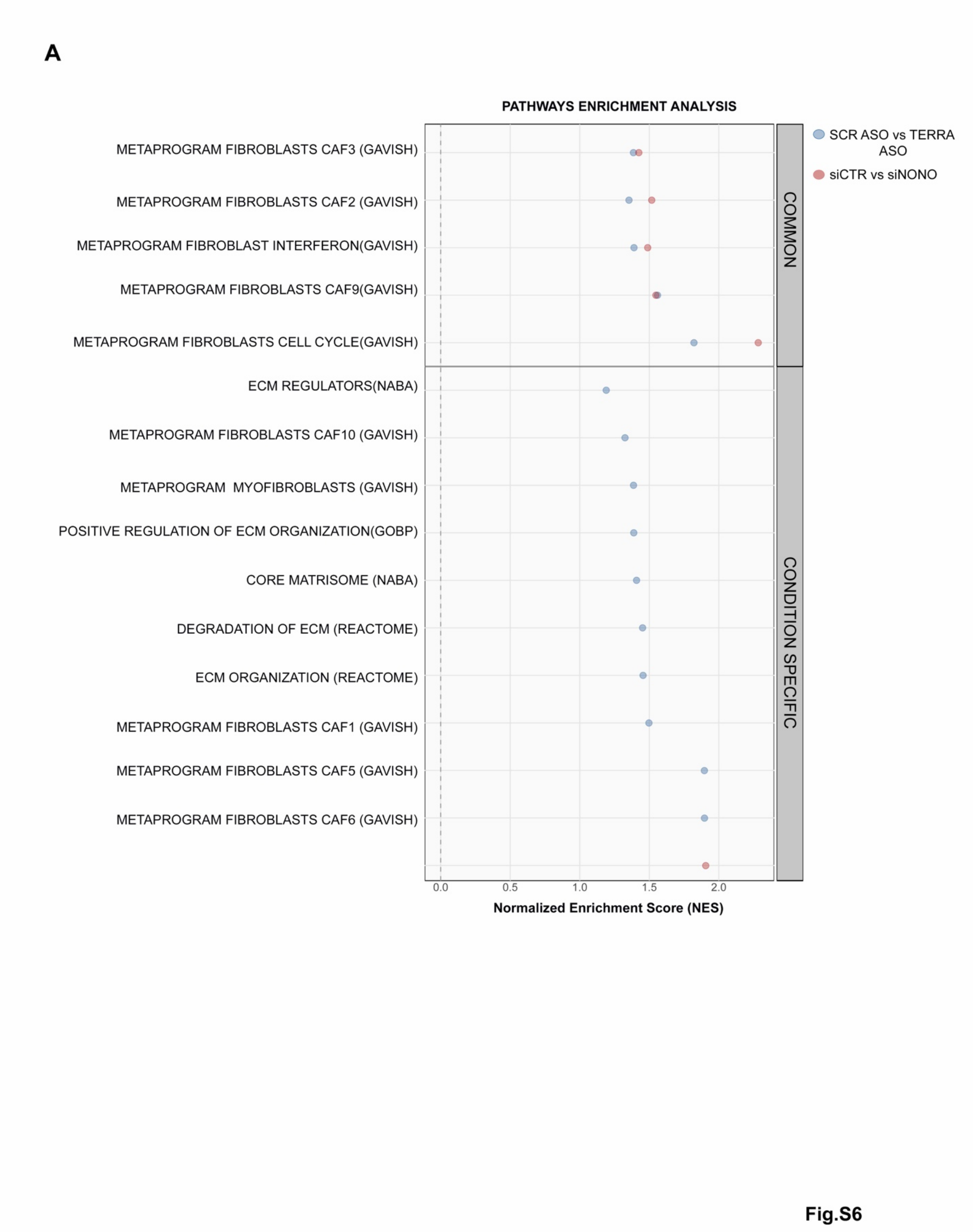


**Fig. S6 Gene Set Enrichment Analysis (GSEA) of CAFs and ECM-related gene signatures in CAFs following TERRA or NONO silencing.**

**(A)**GSEA was performed using curated gene sets from MSigDB (https://www.gsea-msigdb.org/gsea/msigdb/), including: i) extracellular matrix-related signatures (e.g., NABA matrisome, DOI: 10.1074/mcp.M111.014647) ii) fibroblast/CAF metaprograms (Gavish et al.), iii) Reactome degradation of ECM https://www.gsea-msigdb.org/gsea/msigdb/human/geneset/REACTOME_DEGRADATION_OF_THE_EXTRACELLULAR_MATRIX, iv) GOBP_POSITIVE_REGULATION_OF_EXTRACELLULAR_MATRIX_ORGANIZATION (GO:1903055) Transcriptomic profiles were obtained from two CAF strains transfected with control or NONO-targeting siRNAs (siCTR vs. siNONO, red dots), and three CAF strains treated with control or TERRA-targeting antisense oligonucleotides (SCR ASO vs. TERRA ASO, blue dots). The x-axis indicates the Normalized Enrichment Score (NES). Pathways enriched in both conditions are grouped under "Common," while those specific to one condition are listed under "Condition Specific."

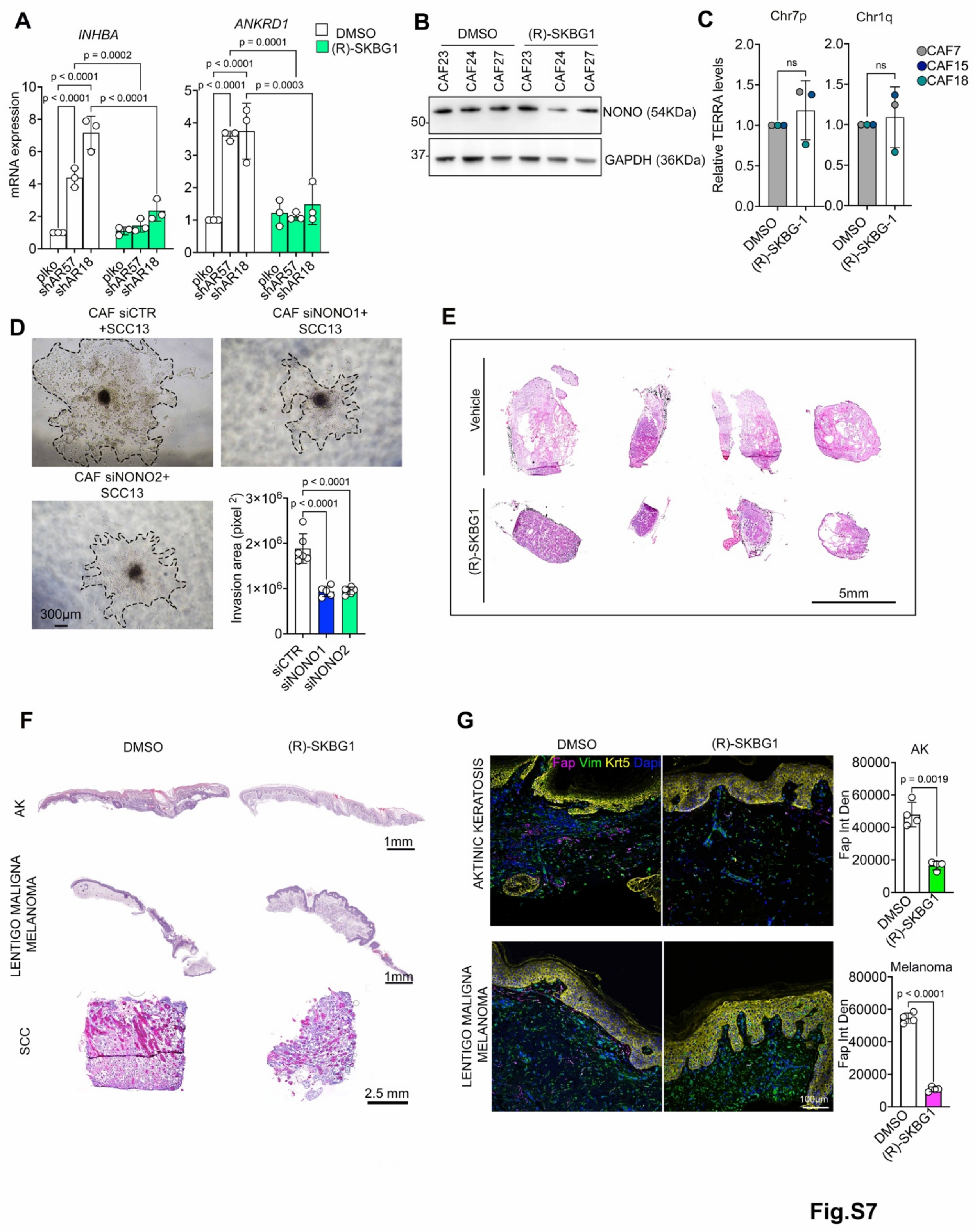


**fig. S7 Pharmacologic and genetic disruption of NONO impairs CAF-mediated tumor progression**

**(A)** Expression of the indicated CAF effector genes in HDFs plus/minus AR silencing, followed by DMSO or (R)-SKBG1 treatment (5μM) for 48 hours, analyzed by RT-qPCR.

Data expressed as fold change relative to DMSO. Data normalized to RPLP0.n=3 biological replicates. Mean ± SD. One-way ANOVA with Šídák's multiple comparisons test.

**(B)** Immunoblot analysis of anti-NONO and GAPDH antibodies of three strains of CAFs, 48 hours after DMSO or (R)-SKBG1 treatment (5 μM).

**(C)** Chromosome-specific (Chr7p, Chr1q) TERRA expression in three CAF strains treated with DMSO or (R)-SKBG1 as described before. Data expressed as fold change relative to DMSO, normalized to GAPDH. Mean ± SD. Paired two-tailed t-test; p-value=0.476 (Cht7p), p=0.7145 (Chr1q).

**(D)** Spheroid invasion assays of SCC13 cells admixed with CAFs transfected with siCTR, or siNONO (1,2). Shown are representative bright field images of spheroids and quantification of the invasion area measured as the difference between the core and the surrounding invaded area delimited with dotted lines. n >5 spheroids per condition. One-way ANOVA with Dunnett's multiple comparisons test.

**(E)** H&E-stained tumor (#2, #3, #4, #5) as described in Fig.7H.

(**F**) H&E staining of the ex vivo actinic keratosis, cSCC and lentigo malignant melanoma treated with DMSO or (R)-SKBG1 (10μM) for 48 hours.

**(G)** Representative immunofluorescence images of actinic keratosis (AK) and lentigo maligna melanoma biopsies treated ex vivo with DMSO or the NONO inhibitor (R)-SKBG1 (10 μM) for 48 h. Sections were stained for FAP (Fap; magenta), vimentin (Vim; green), keratin 5 (Krt5; yellow), and DAPI (blue). Lower panels: quantification of FAP immunofluorescence signal intensity (integrated density) in AK and melanoma lesions. Each point represents one field. Mean ± SD; Unpaired t test with Welch's correction.

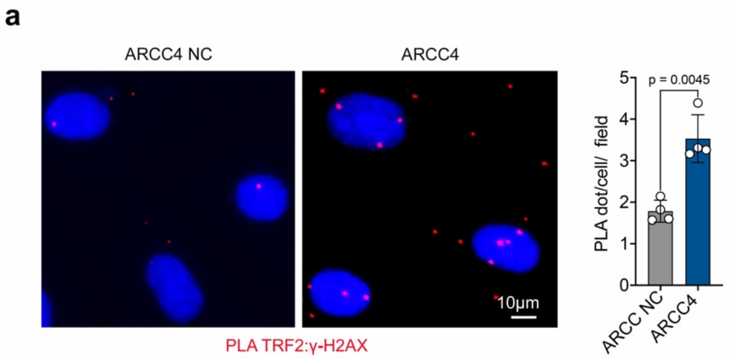


**Fig. S8 AR degradation promotes telomeric DNA damage in HDFs.**

**(A)** PLA assays with antibodies against TRF2 and γ-H2AX in HDFs with ARCC4 NC or ARCC4 treatment (as described in Fig. 2c). Shown are representative images and quantification of the number of PLA puncta per cell per field. Data represent pooled fields from 3 independent experiments (n>4 fields total, counting > 200 cells per field). Mean ± SD, Unpaired t-test with Welch's correction.
